## Supplementary material for "Glycoengineered keratinocyte library reveals essential functions of specific glycans for all stages of HSV-1 infection": Table S1

Table S1. Knock out indel sequencing and sgRNA/ZFN targeting sequences

| Gene | Clone | Sequence^[1]^ | gRNA/ZFN (5’-3’) | Indel | First affected a.a. |
| --- | --- | --- | --- | --- | --- |
| N-Glycan |  |  |  |  |  |
| *MGAT1* KO | REF | GATtcgcctggcccaagacgccgAGG | TCGCCTGGCCCAAGACGCCG |  |  |
|  | 1E7_A1 | GATtcgcctggcccaag----cgCGG |  | -4 | 60 |
|  | 1E7_A2 | GATtcgcctggcccaagacg-cgAGG |  | -1 | 61 |
| *MGAT4A* KO | REF | aatgagatgaggctccgcaaTGGAAC | AATGAGATGAGGCTCCGCAA |  |  |
|  | 3C10_A1 | aatgagatgaggctcc--aaTGGAAC |  | -2 | 4 |
|  | 3C10_A2 | aatgagatgaggctccgca-TGGAAC |  | -1 | 5 |
| *MGAT4B* KO | REF | TTG…GCAggagagcctcaagc-gctccaAGGAGCTCAACCTGGTGCTGG | GGAGAGCCTCAAGCGCTCCA |  |  |
|  | 3F8_A1 | TTG…GCAggagagcctcaagcTgctccaAGGAGCTCAACCTGGTGCTGG |  | +1 | 60 |
|  | 3F8_A2 | TTG…------------------------------------------TGG |  | -96 | 36 |
| *MGAT5+4B*  KO | 2B5 A1 (in M*GAT5* KO 1F12) | TTG…GCAgga--------------------GAGCTCAACCTGGTGCTGG |  | -19 | 56 |
| *MGAT5* KO | REF | GGGCCAtac-------gctggagtcatgacagcTTAT | GCTGTCATGACTCCAGCGTA |  |  |
|  | 1F12_A1 | GGGCCAtacA------gctggagtcatgacagcTTAT |  | +1 | 74 |
|  | 1F12_A2 | GGGCCACATGACATAGgctggagtcatgacagcTTAT |  | +7 | 73 |
| O-GalNAc |  |  |  |  |  |
| *C1GALT1C1* KO^[2]^ | REF | ATCACCCCAACCAGGTAGTagaaggctGTTGTTCAGATAT | AGAAGGCT (ZFN cut site) |  |  |
|  | D5_A1 | ATCACCCCAACCAGGTA----------GTTGTTCAGATAT |  | -10 | 267 |
| *C1GALT1* KO | REF | GCCCAgcgttgtaacaaagtgttgtTTATGAGTTCAGAA | ACAACACTTTGTTACAACGC |  |  |
|  | 2E5_A1 | GCCCAG--TTGTAACAAAGTGTTGTTTATGAGTTCAGAA |  | -2 | 114 |
|  | 2E5_A2 | GCCCAGCG-----------TGTTGTTTATGAGTTCAGAA |  | -11 | 115 |
| *GCNT1* KO | REF | AATCACCTtctccgttttaaggattcatCAA | ATGAATCCTTAAAACGGAGA |  |  |
|  | 2E7_A1 | AAT-------------ttaaggattcatCAA |  | -13 | 27 |
|  | 2E7_A2 | AATCACCTtc--cgttAATGggattcatCAA |  | -2 | 30 |
| *ST3GAL1* KO | REF | TGGTGCCCttc-aagaccatcgacttggaGTGGG | TCCAAGTCGATGGTCTTGAA |  |  |
|  | 2B2_A1 | TGGTGCCCttcAaagaccatcgacttggaGTGGG |  | +1 | 210 |
| GAG |  |  |  |  |  |
| *B4GALT7* KO | REF | TGCACGACGTtgacctgctccctctcaa-cgAGGAGCTGGAC | TGACCTGCTCCCTCTCAACG |  |  |
|  | 3D9_A1 | TGCACGACGTtgacctgctccctctcaaAcgAGGAGCTGGAC |  | +1 | 170 |
|  | 3D9_A2 | TGCACGACGTtgacctgctccctctc--------------AC |  | -13 | 170 |
| GSL |  |  |  |  |  |
| *B4GALT5* KO | REF | AGCCTTCTGATTGCat-------------gccTCGGTGGAAG | ATGCC (ZFN cut site) |  |  |
|  | D11_A1 | AGCCTTCTGATTGCATGTAGAATTCCCTAGCCTCGGTGGAAG |  | +13 |  |
|  |  |  |  |  | 160 |
| *ST3GAL5* KO | REF | GTTatttgagcacaggtata-gcgTGGACTTAC | ATTTGAGCACAGGTATAGCG |  |  |
|  | 1C5_A1 | GTTatttgagcacaggtataCgcgTGGACTTAC |  | +1 | 138 |
|  | 1C5_A2 | GTTatttgagcGTAA---------TGGACTTAC |  | -8 | 135 |

1. Sequences are shown in the orientation of the gene, with sgRNA recognition sequences (direct or complementary)/ZFN cut sites in lower case and PAM sequences underlined
2. Radhakrishnan *et al*., 2014 (DOI: 10.1073/pnas.1406619111)
