## Supplementary material for "Glycoengineered keratinocyte library reveals essential functions of specific glycans for all stages of HSV-1 infection": Table S2

Table S2. Sequences of mutagenic primers. Altered nucleotides are shown in red.

| Name | 5’ to 3’ | Length |
| --- | --- | --- |
| gB_T169A | GTACTACAAAGACGTCGCCGTTTCGCAGGTGTG | 33 |
| gB_T267A | CCACCGGTACGGGGCGACGGTAAACTG | 27 |
| gB_T268A | ACCGGTACGGGACGGCGGTAAACTGCATC | 29 |
| gB_T267A_T268A | TCCACCGGTACGGGGCGGCGGTAAACTGCATC | 32 |
| gB_T690A | CCTGGAGGTGTACGCCCGCCACGAGATC | 28 |
| gB_T703A | CCTGCTGGACTACGCGGAGGTCCAGCG | 27 |
| gD_S33A | CCTTGGCGGATGCCGCTCTCAAGATGGCC | 29 |
| gD_T255A | CGAGAACCAGCGCGCCGTCGCCGTATA | 27 |
| gD_S260A | ACCGTCGCCGTATACGCCTTGAAGATCGCCGG | 32 |
| gD_T255A_S260A | CGAGAACCAGCGCGCCGTCGCCGTATACGCCTTGAAGATCGCCGG | 45 |
