## Supplementary figures and images for "Glycoengineered keratinocyte library reveals essential functions of specific glycans for all stages of HSV-1 infection"

### Figure S1

Figure S1. N-glycoprofiling

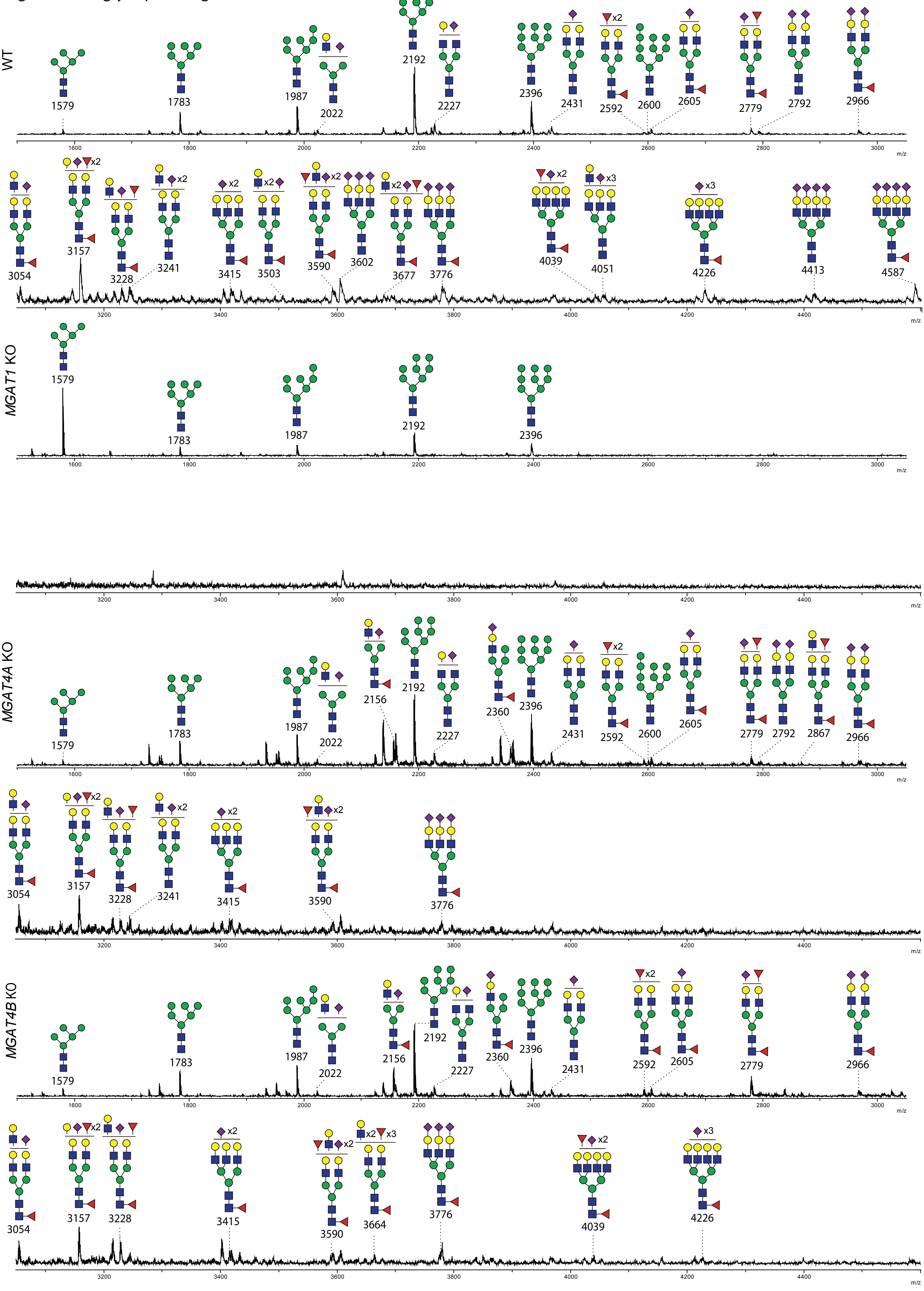

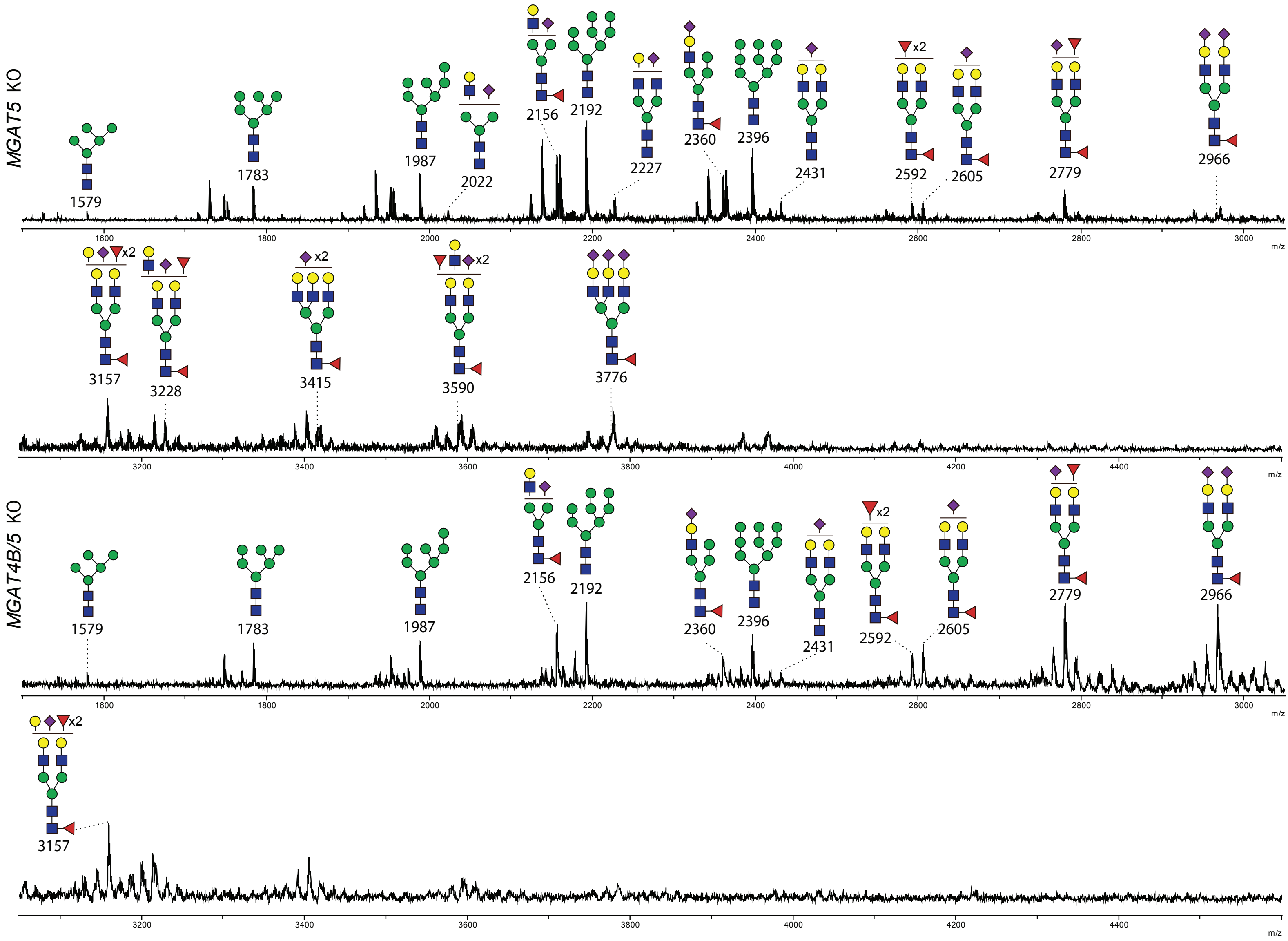

### Figure S2

Figure S2. CORA O-glycoprofiling

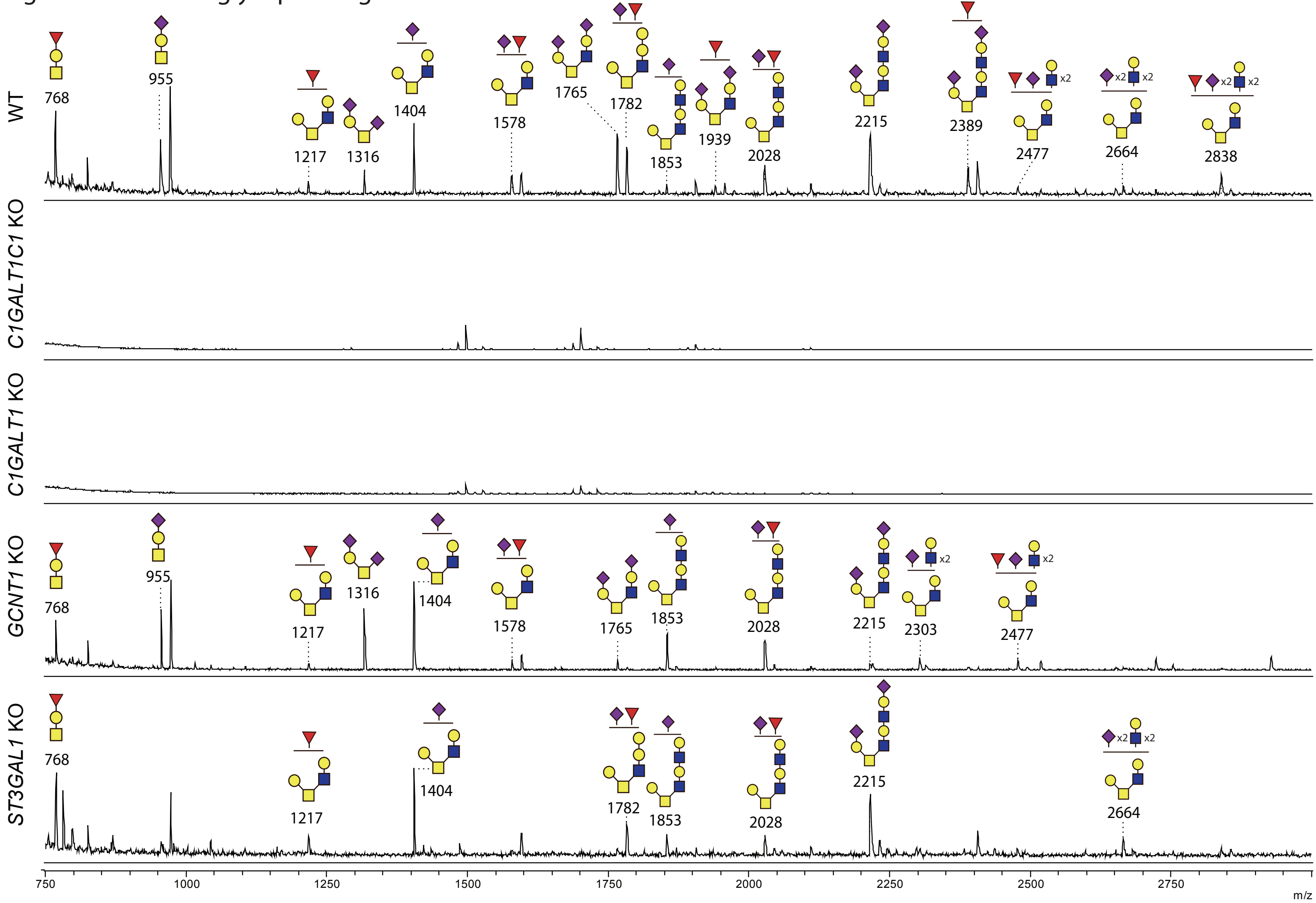
