## Supplementary material for "Glycoengineered keratinocyte library reveals essential functions of specific glycans for all stages of HSV-1 infection": Figure S3

**Figure S3. Sequencing of DNA constructs.** Multiple sequence alignment for the DNA regions corresponding to the positions of introduced single amino acid mutations in HSV-1 gB and gD expression constructs. Asterisks indicate identical nucleotides.

```
gB Tyr165-Val173          TyrLysAspValThrValSerGlnVal
                           Y  K  D  V  T  V  S  Q  V
gB_WT                     TACAAAGACGTCACCGTTTCGCAGGTG   27
gB_T169A                  TACAAAGACGTCGCCGTTTCGCAGGTG   27
gB_T267A                  TACAAAGACGTCACCGTTTCGCAGGTG   27
gB_T268A                  TACAAAGACGTCACCGTTTCGCAGGTG   27
gB_T267A_T268A            TACAAAGACGTCACCGTTTCGCAGGTG   27
gB_T690A                  TACAAAGACGTCACCGTTTCGCAGGTG   27
gB_T703A                  TACAAAGACGTCACCGTTTCGCAGGTG   27
gB_T690A_T703A            TACAAAGACGTCACCGTTTCGCAGGTG   27
gB_T169A_T267A_T268A      TACAAAGACGTCGCCGTTTCGCAGGTG   27
                          ************ **************

gB His263-Ile272          HisArgTyrGlyThrThrValAsnCysIle
                           H  R  Y  G  T  T  V  N  C  I
gB_WT                     CACCGGTACGGGACGACGGTAAACTGCATC   30
gB_T169A                  CACCGGTACGGGACGACGGTAAACTGCATC   30
gB_T267A                  CACCGGTACGGGGCGACGGTAAACTGCATC   30
gB_T268A                  CACCGGTACGGGACGGCGGTAAACTGCATC   30
gB_T267A_T268A            CACCGGTACGGGGCGGCGGTAAACTGCATC   30
gB_T690A                  CACCGGTACGGGACGACGGTAAACTGCATC   30
gB_T703A                  CACCGGTACGGGACGACGGTAAACTGCATC   30
gB_T690A_T703A            CACCGGTACGGGACGACGGTAAACTGCATC   30
gB_T169A_T267A_T268A      CACCGGTACGGGGCGGCGGTAAACTGCATC   30
                          ************ ** **************

gB Glu687-Gln706          GluValTyrThrArgHisGluIleLysAspSerGlyLeuLeuAspTyrThrGluValGln
                           E  V  Y  T  R  H  E  I  K  D  S  G  L  L  D  Y  T  E  V  Q
gB_WT                     GAGGTGTACACCCGCCACGAGATCAAGGACAGCGGCCTGCTGGACTACACGGAGGTCCAG   60
gB_T169A                  GAGGTGTACACCCGCCACGAGATCAAGGACAGCGGCCTGCTGGACTACACGGAGGTCCAG   60
gB_T267A                  GAGGTGTACACCCGCCACGAGATCAAGGACAGCGGCCTGCTGGACTACACGGAGGTCCAG   60
gB_T268A                  GAGGTGTACACCCGCCACGAGATCAAGGACAGCGGCCTGCTGGACTACACGGAGGTCCAG   60
gB_T267A_T268A            GAGGTGTACACCCGCCACGAGATCAAGGACAGCGGCCTGCTGGACTACACGGAGGTCCAG   60
gB_T690A                  GAGGTGTACGCCCGCCACGAGATCAAGGACAGCGGCCTGCTGGACTACACGGAGGTCCAG   60
gB_T703A                  GAGGTGTACACCCGCCACGAGATCAAGGACAGCGGCCTGCTGGACTACGCGGAGGTCCAG   60
gB_T690A_T703A            GAGGTGTACGCCCGCCACGAGATCAAGGACAGCGGCCTGCTGGACTACGCGGAGGTCCAG   60
gB_T169A_T267A_T268A      GAGGTGTACACCCGCCACGAGATCAAGGACAGCGGCCTGCTGGACTACACGGAGGTCCAG   60
                          ********* ************************************** ***********

gD Leu29-Ala37            LeuAlaAspAlaSerLeuLysMetAla
                           L  A  D  A  S  L  K  M  A
gD_WT                     TTGGCGGATGCCTCTCTCAAGATGGCC   27
gD_S33A                   TTGGCGGATGCCGCTCTCAAGATGGCC   27
gD_T255A                  TTGGCGGATGCCTCTCTCAAGATGGCC   27
gD_S260A                  TTGGCGGATGCCTCTCTCAAGATGGCC   27
gD_T255A_S260A            TTGGCGGATGCCTCTCTCAAGATGGCC   27
gD_S33A_T255A_S260A       TTGGCGGATGCCGCTCTCAAGATGGCC   27
                          ************ **************

gD Glu251-Ala264          GluAsnGlnArgThrValAlaValTyrSerLeuLysIleAla
                           E  N  Q  R  T  V  A  V  Y  S  L  K  I  A
gD_WT                     GAGAACCAGCGCACCGTCGCCGTATACAGCTTGAAGATCGCC   42
gD_S33A                   GAGAACCAGCGCACCGTCGCCGTATACAGCTTGAAGATCGCC   42
gD_T255A                  GAGAACCAGCGCGCCGTCGCCGTATACAGCTTGAAGATCGCC   42
gD_S260A                  GAGAACCAGCGCACCGTCGCCGTATACGCCTTGAAGATCGCC   42
gD_T255A_S260A            GAGAACCAGCGCGCCGTCGCCGTATACGCCTTGAAGATCGCC   42
gD_S33A_T255A_S260A       GAGAACCAGCGCGCCGTCGCCGTATACGCCTTGAAGATCGCC   42
                          ************ **************  *************
```
