## Supplementary material for "Glycoengineered keratinocyte library reveals essential functions of specific glycans for all stages of HSV-1 infection": Figure S4

**Figure S4. Section panels of HSV-1-infected glycoengineered organotypic skin.** 36 hours post-infection the tissues were fixed in formalin and two independent series of 10 consecutive sections with 30 µm intervals were stained with goat anti-HSV-1 FITC pAb followed by imaging with a fluorescence-equipped slide scanner.

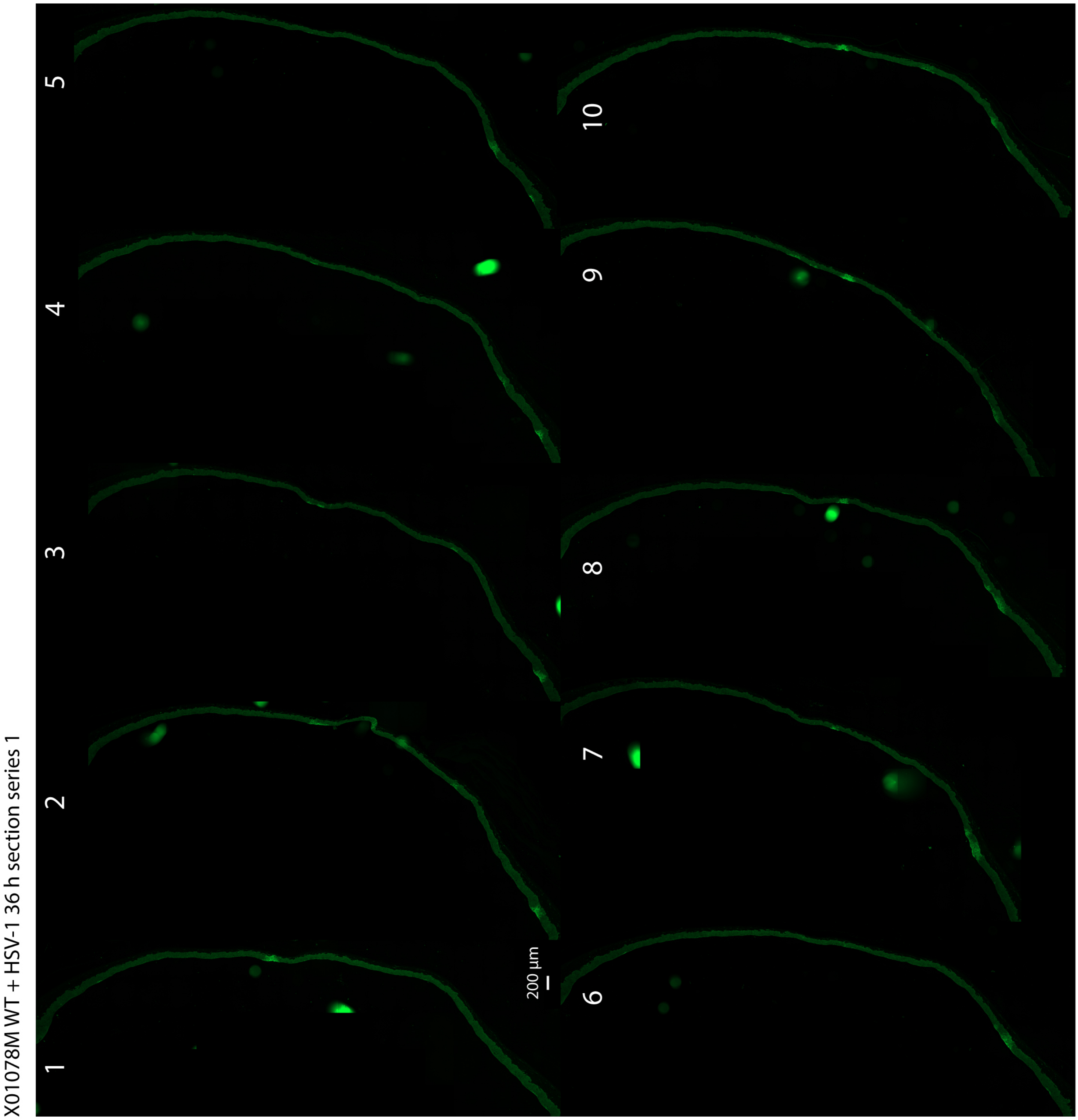

X01078M WT + HSV-1 36 h section series 2

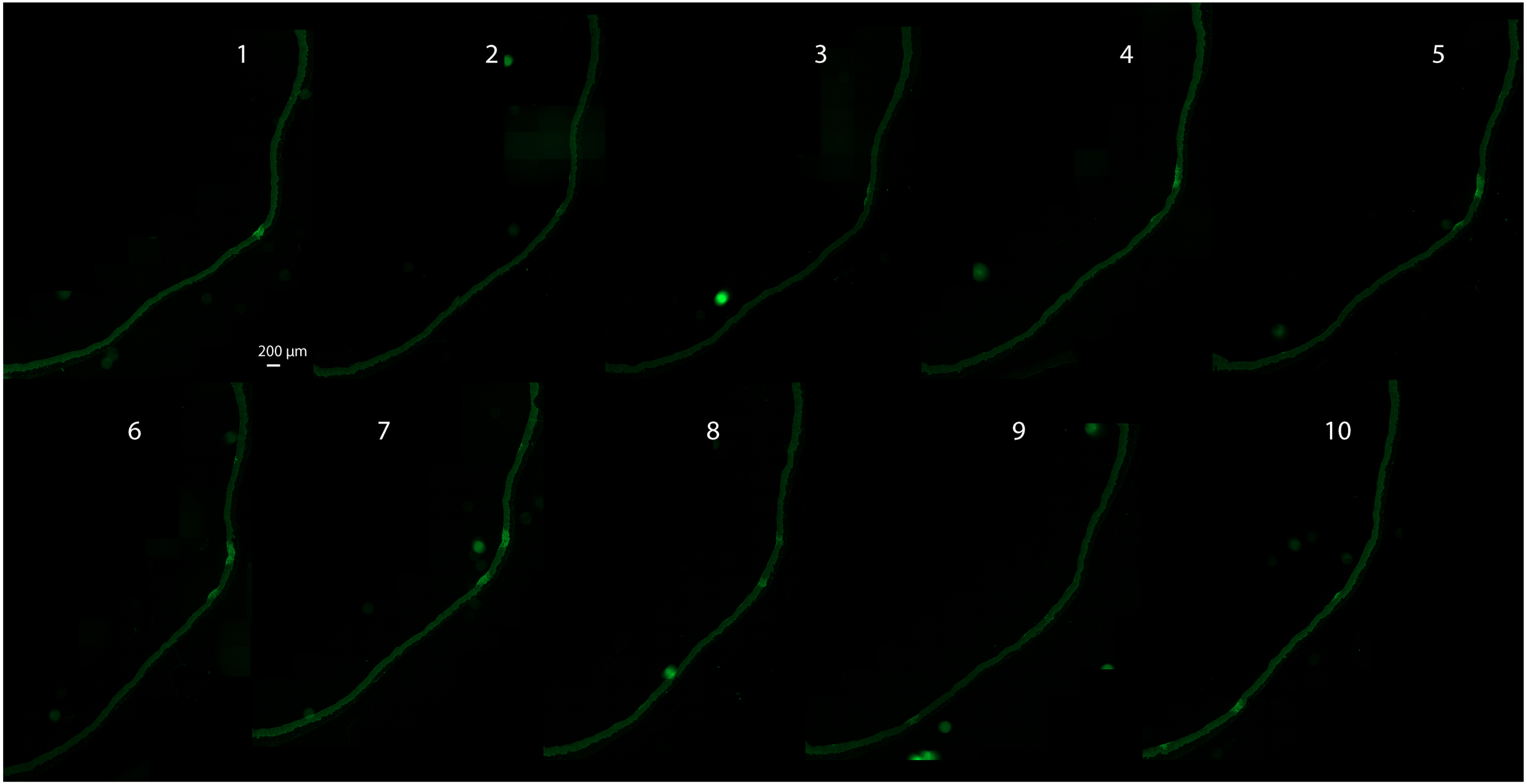

X01078O *MGAT1* KO 1E7 + HSV-1 36 h section series 1

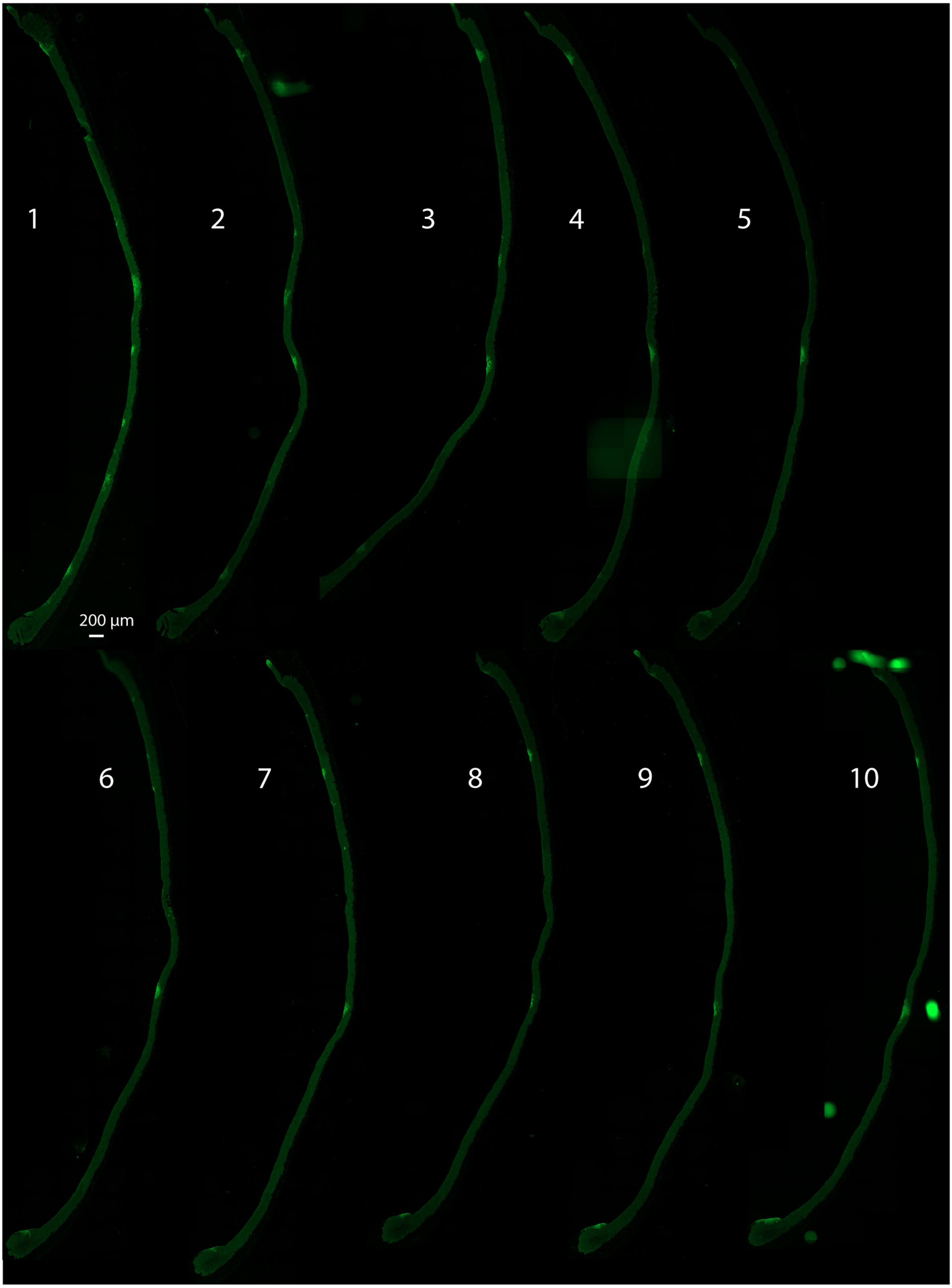

X01078O *MGAT1* KO 1E7 + HSV-1 36 h section series 2

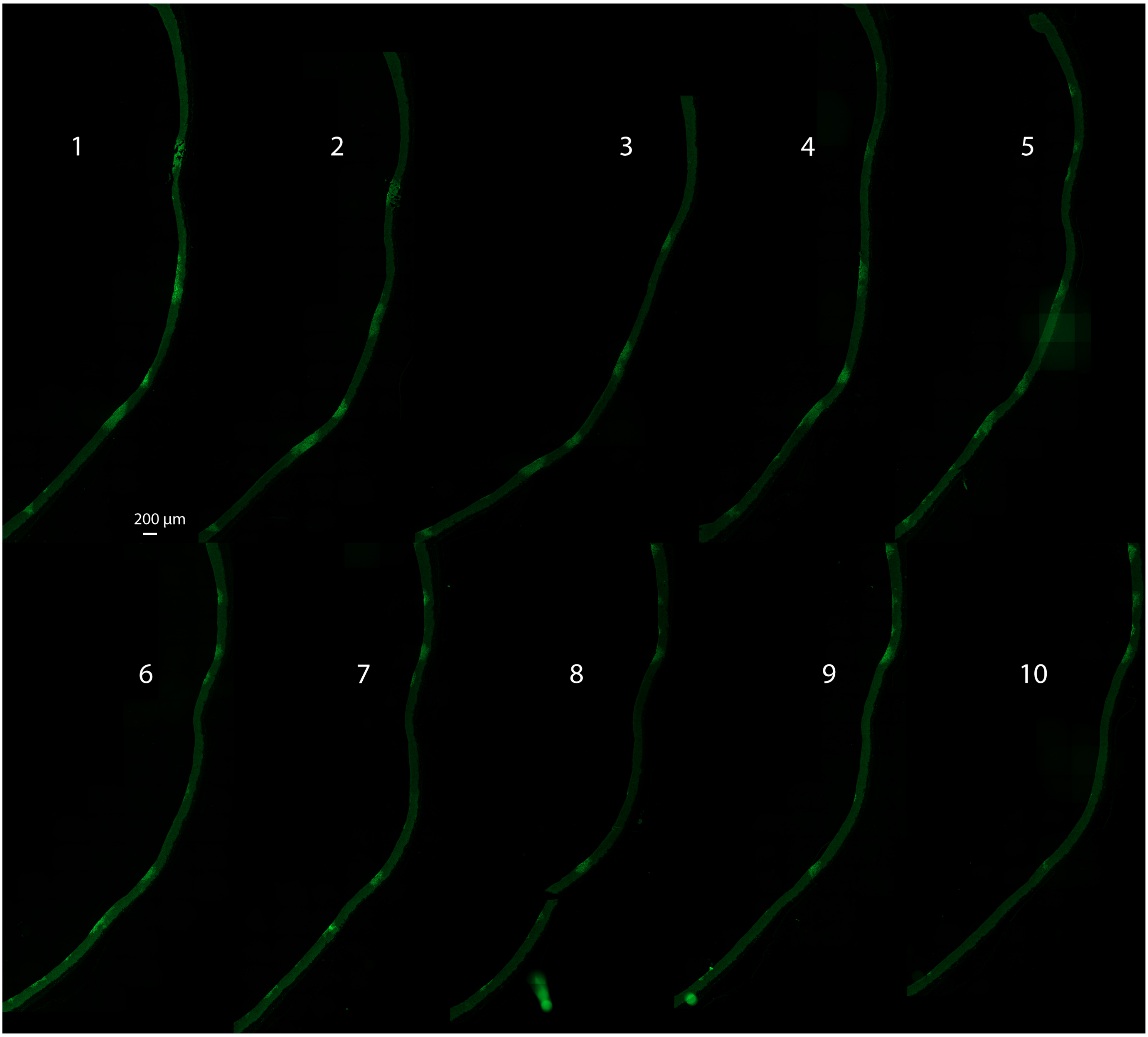

X01078P *MGAT4A* KO 3C10 + HSV-1 36 h section series 1

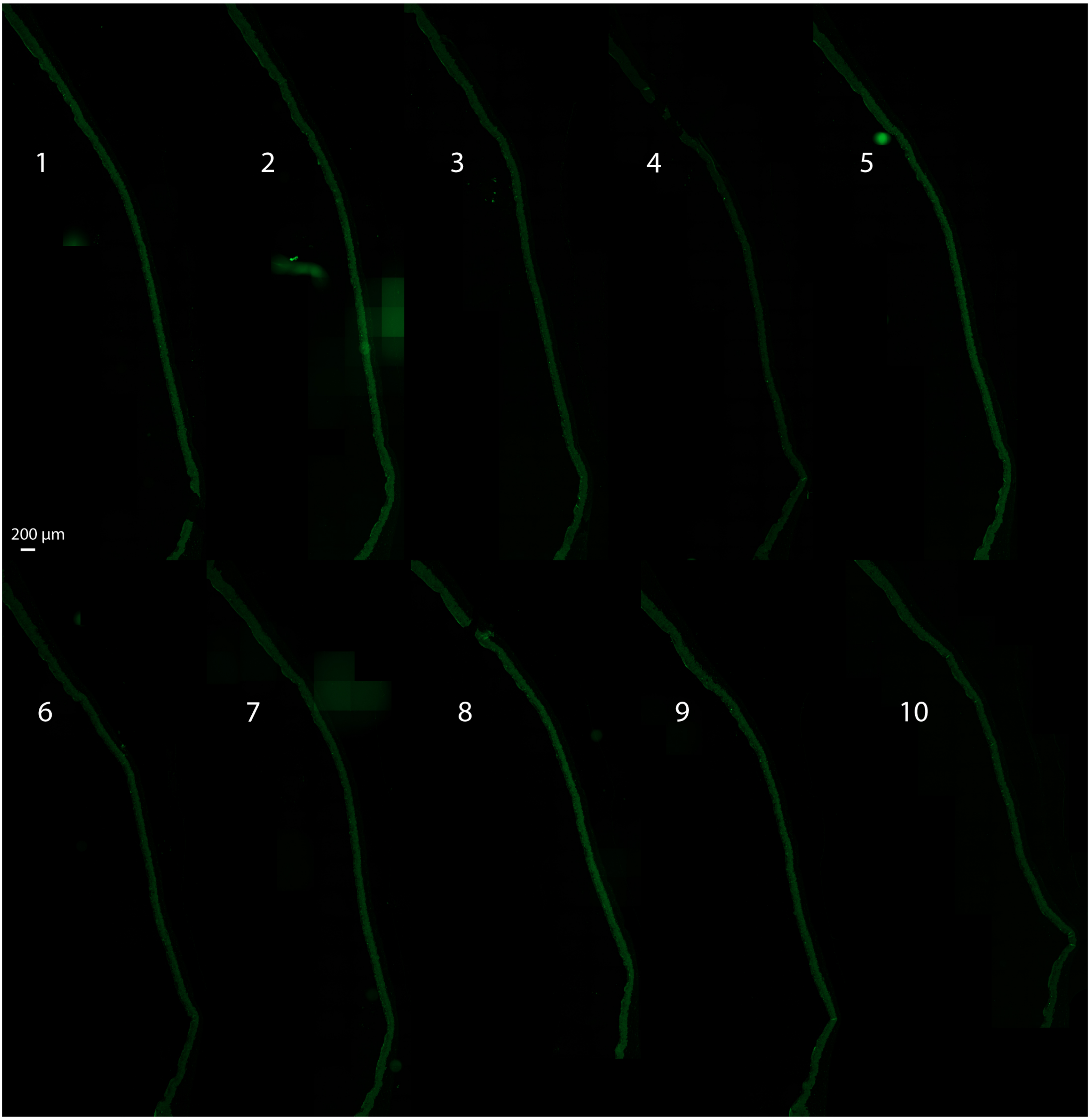

X01078P *MGAT4A* KO 3C10 + HSV-1 36 h section series 2

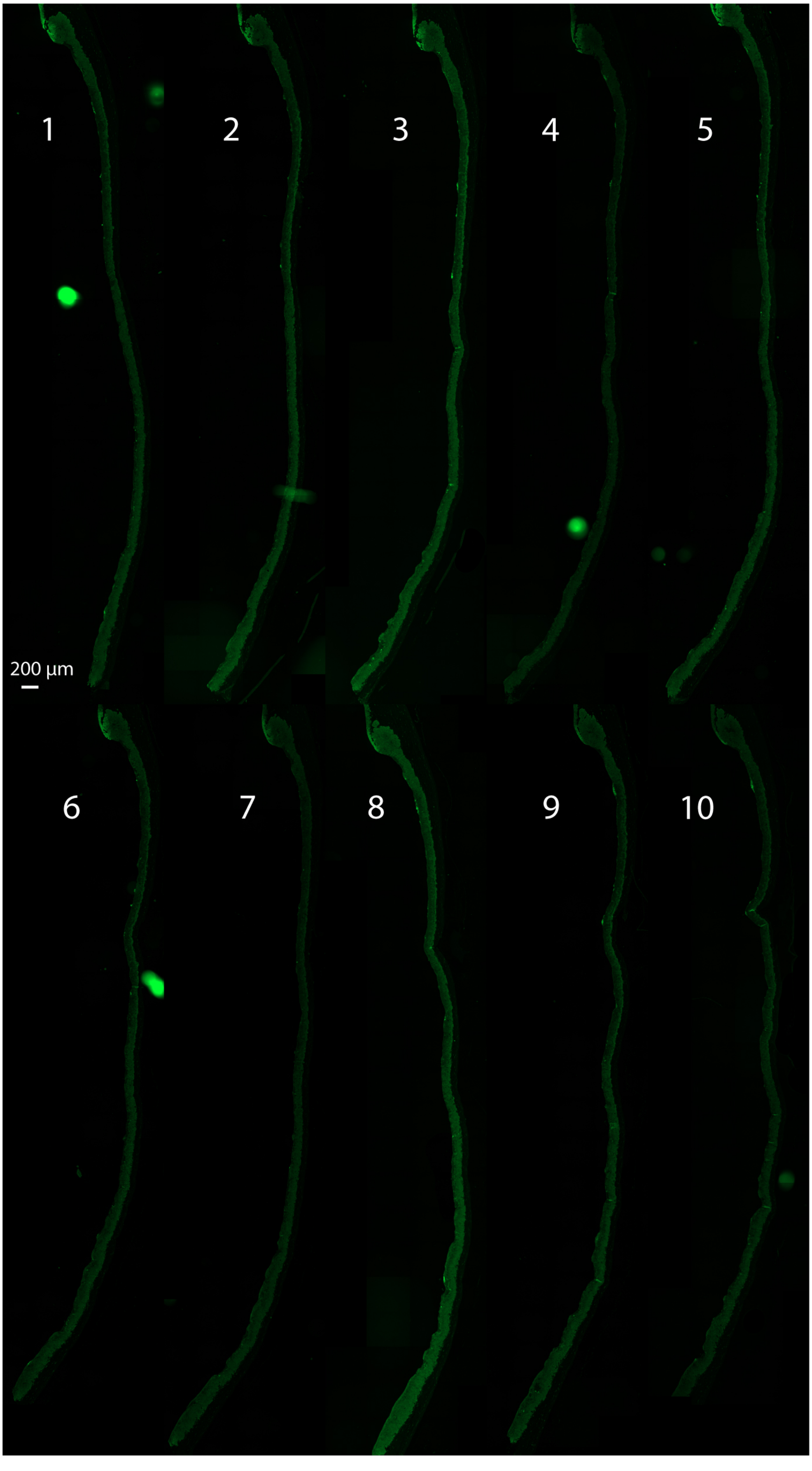

X01105L *MGAT4B* KO 3F8 + HSV-1 36 h section series 1

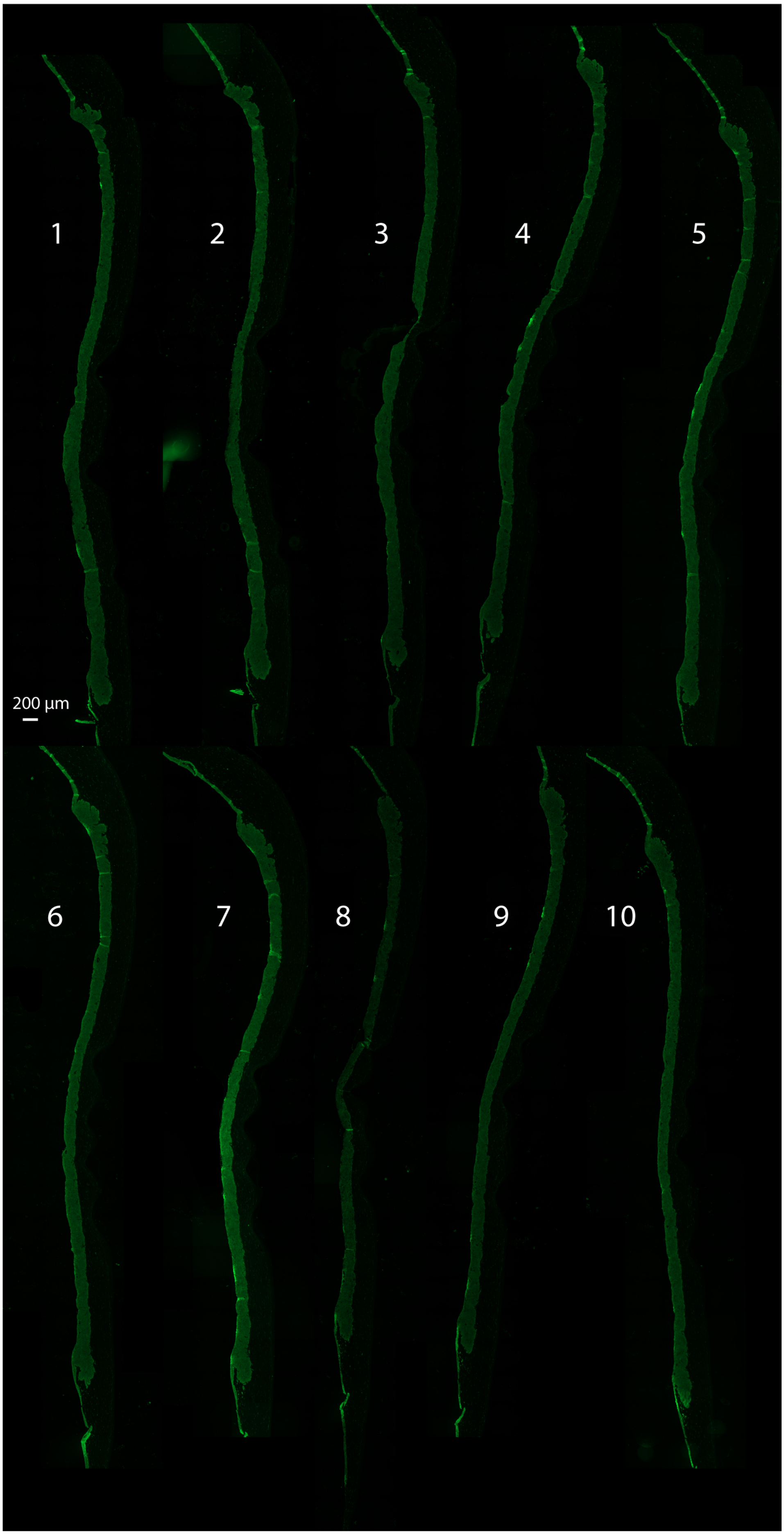

X01105L *MGAT4B* KO 3F8 + HSV-1 36 h section series 2

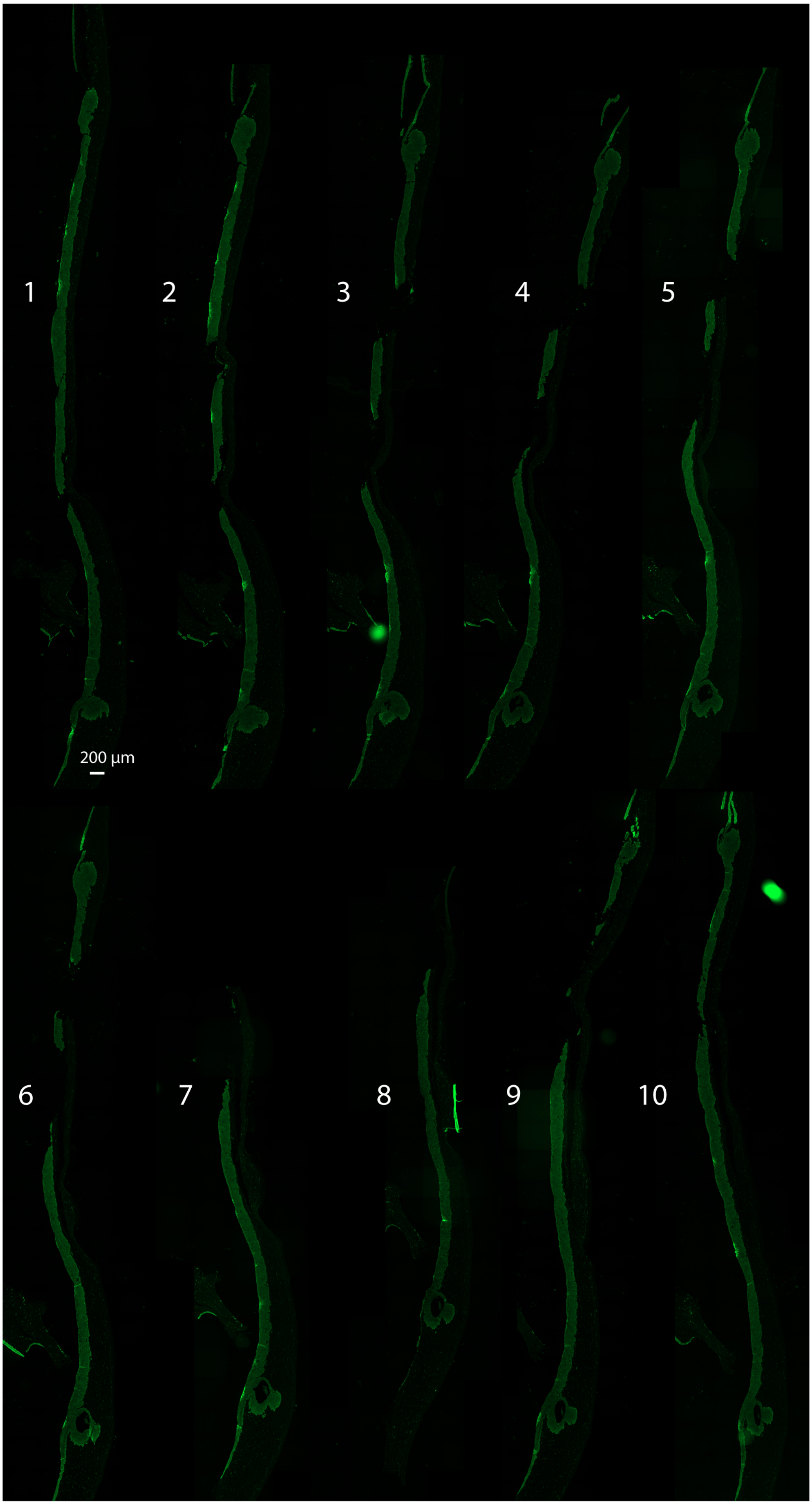

X01078Q *MGAT5* KO 1F12 + HSV-1 36 h section series 1

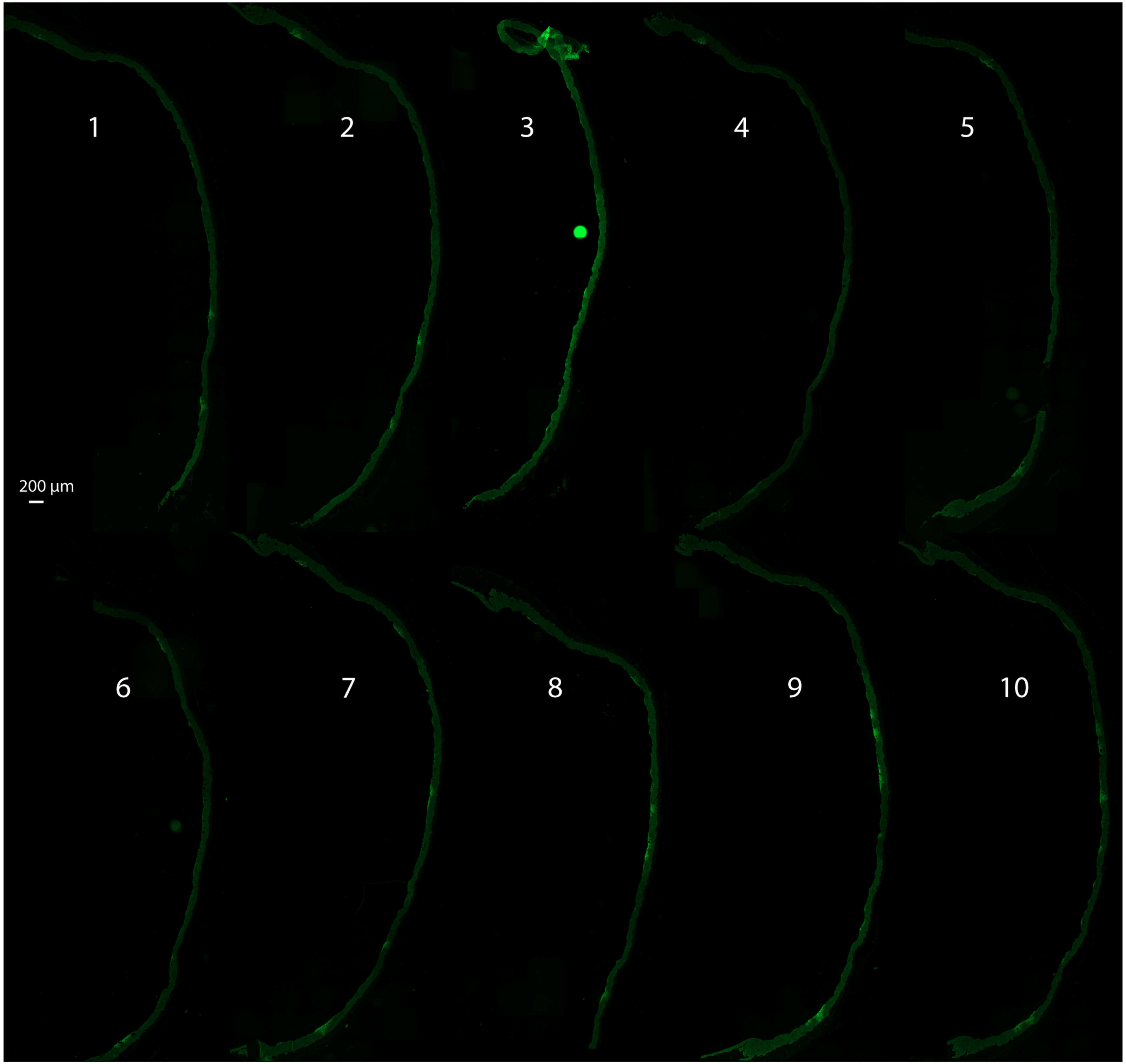

X01078Q *MGAT5* KO 1F12 + HSV-1 36 h section series 2

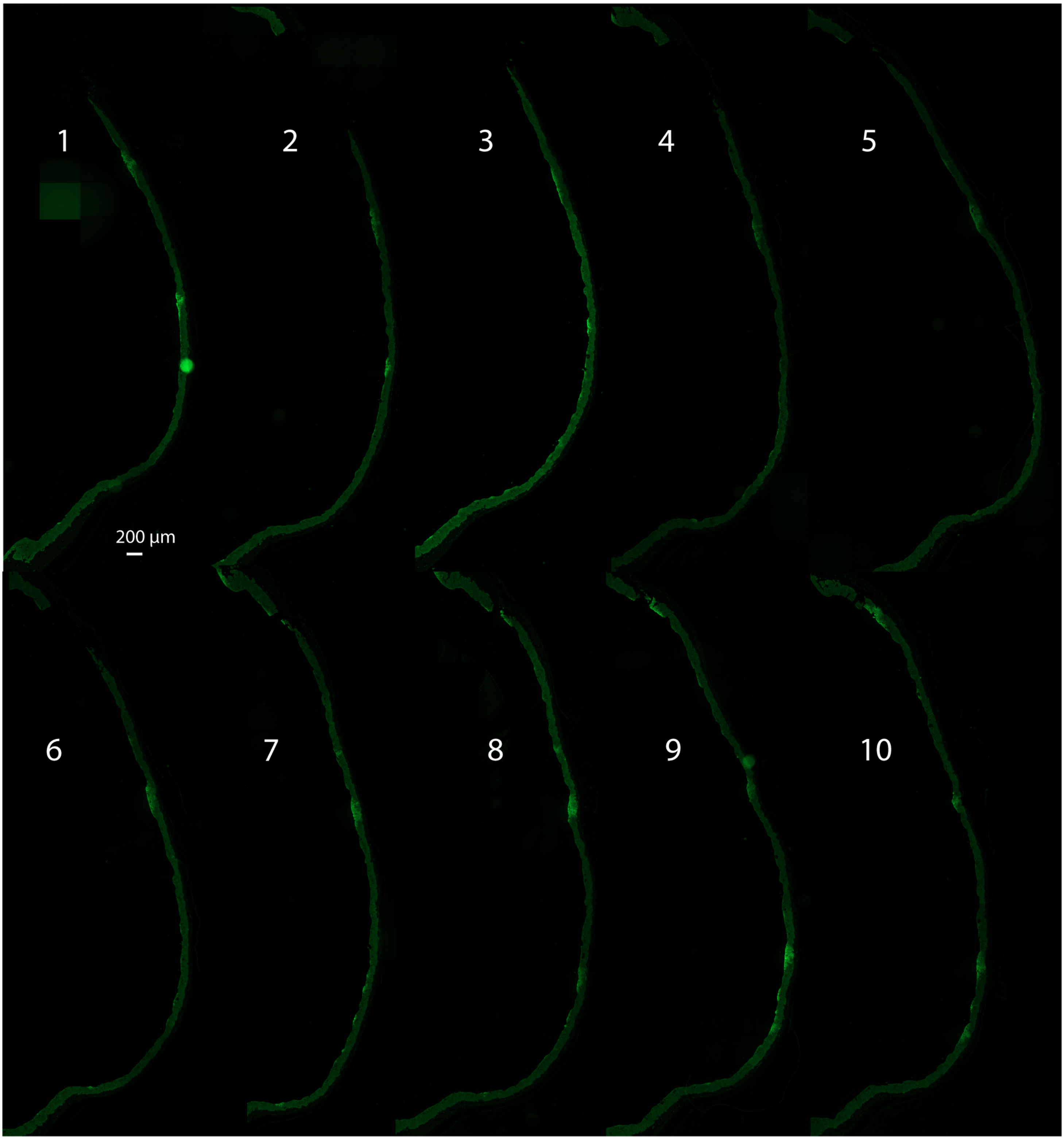

X01105R *MGAT5+4B* KO 2B5 + HSV-1 36 h section series 1

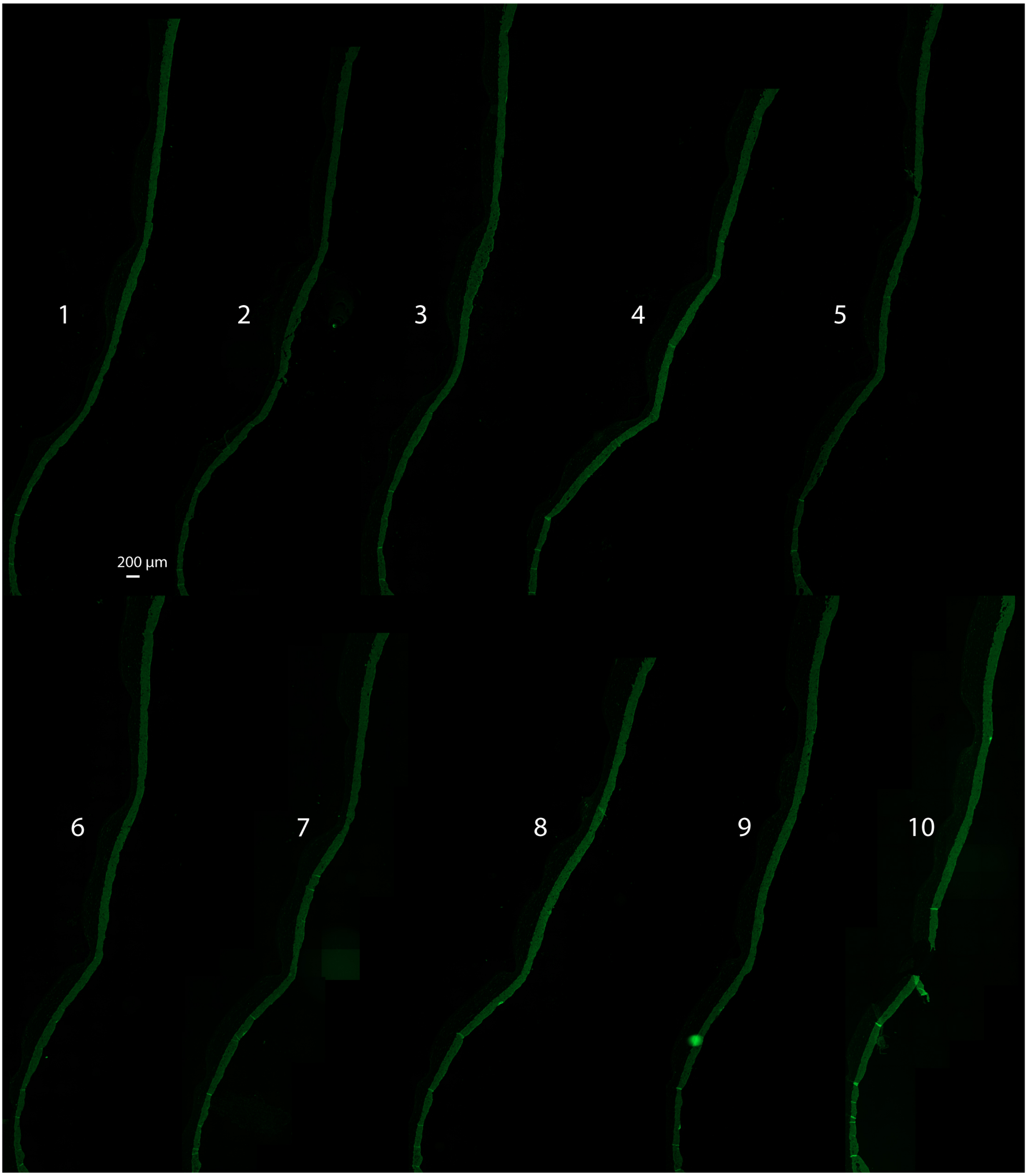

X01105R *MGAT5+4B* KO 2B5 + HSV-1 36 h section series 2

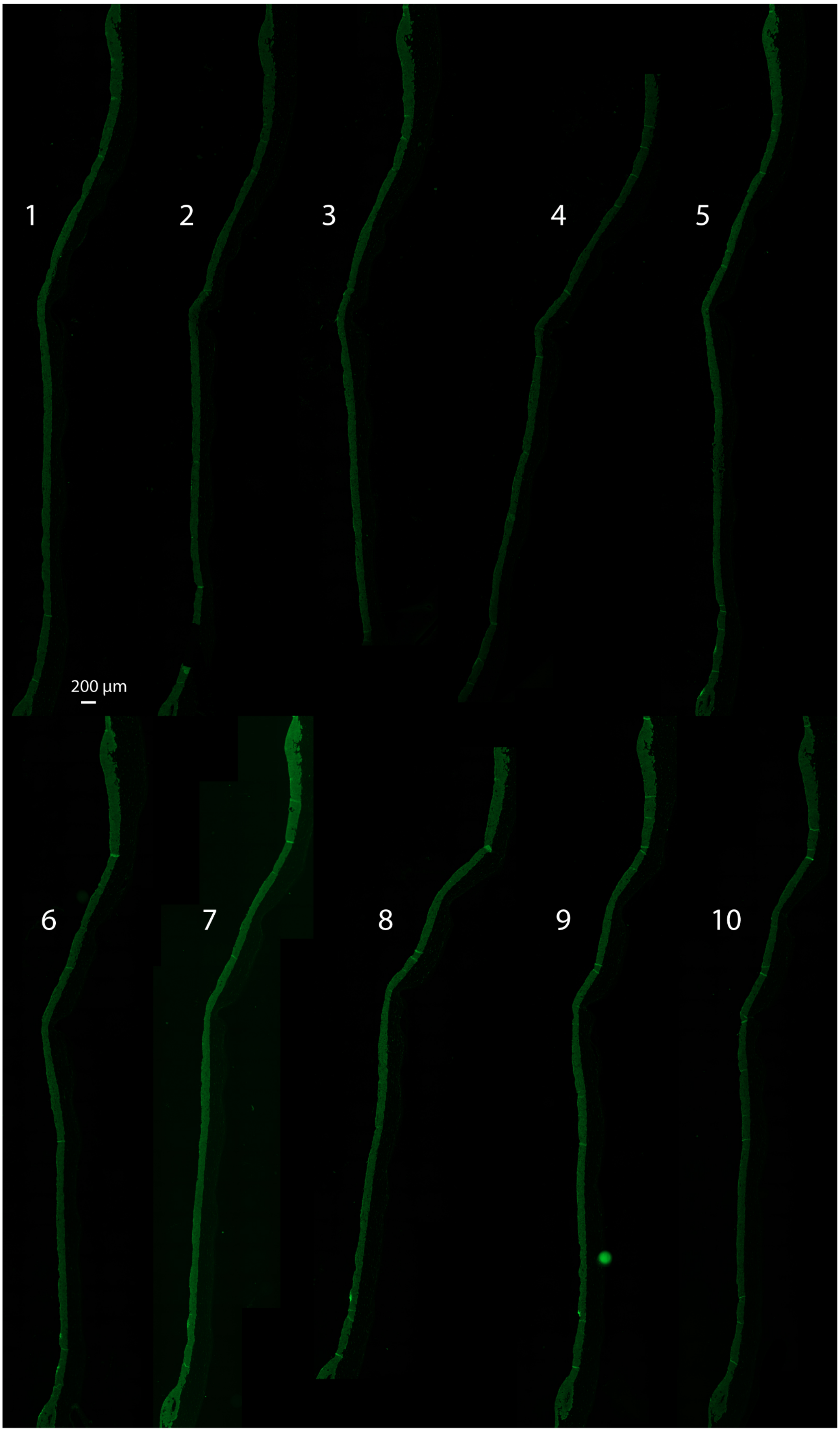

X01078N *C1GALT1C1* KO D5 + HSV-1 36 h section series 1

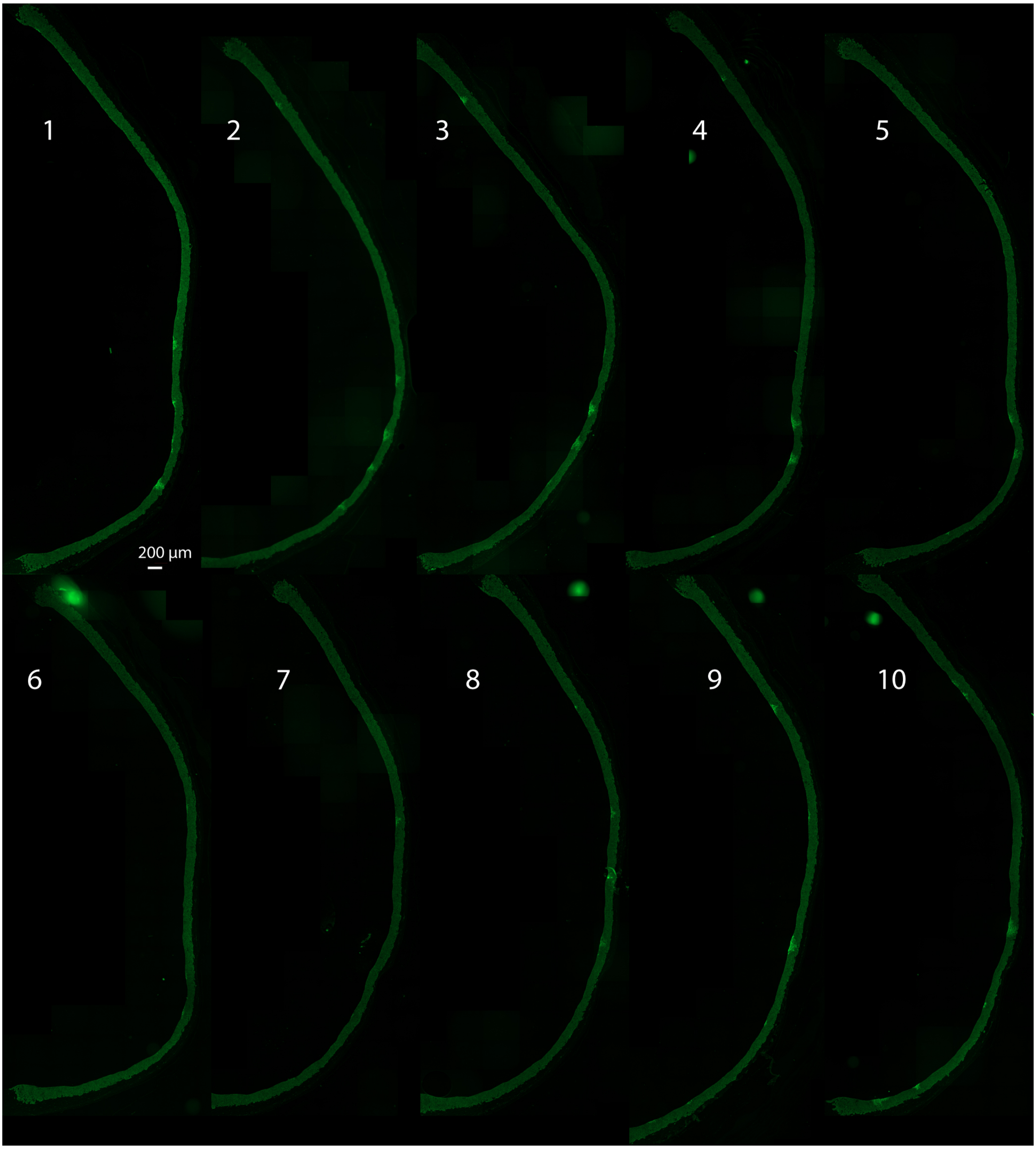

X01078N *C1GALT1C1* KO D5 + HSV-1 36 h section series 2

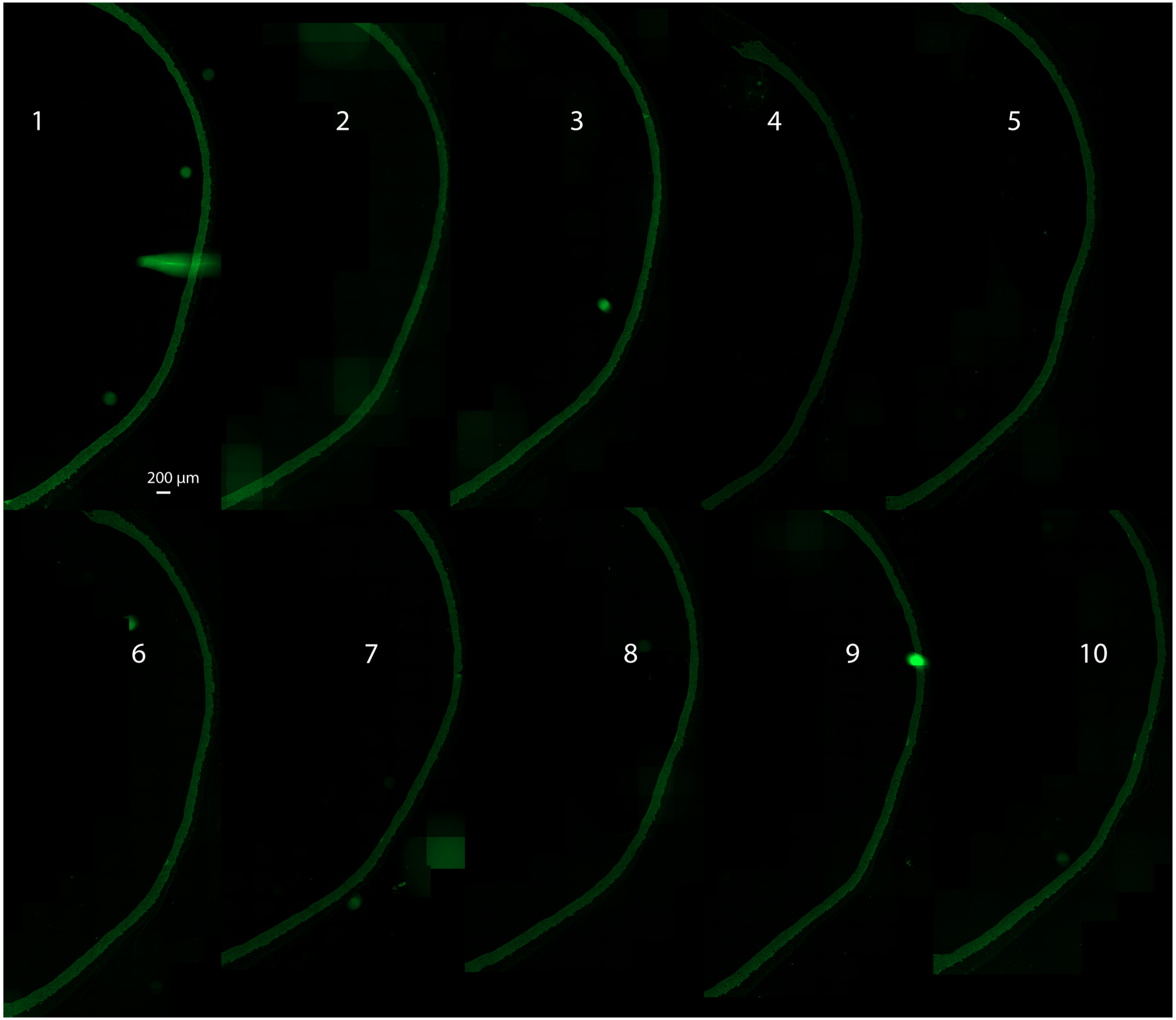

X01105I *C1GALT1* KO 2E5 + HSV-1 36 h section series 1

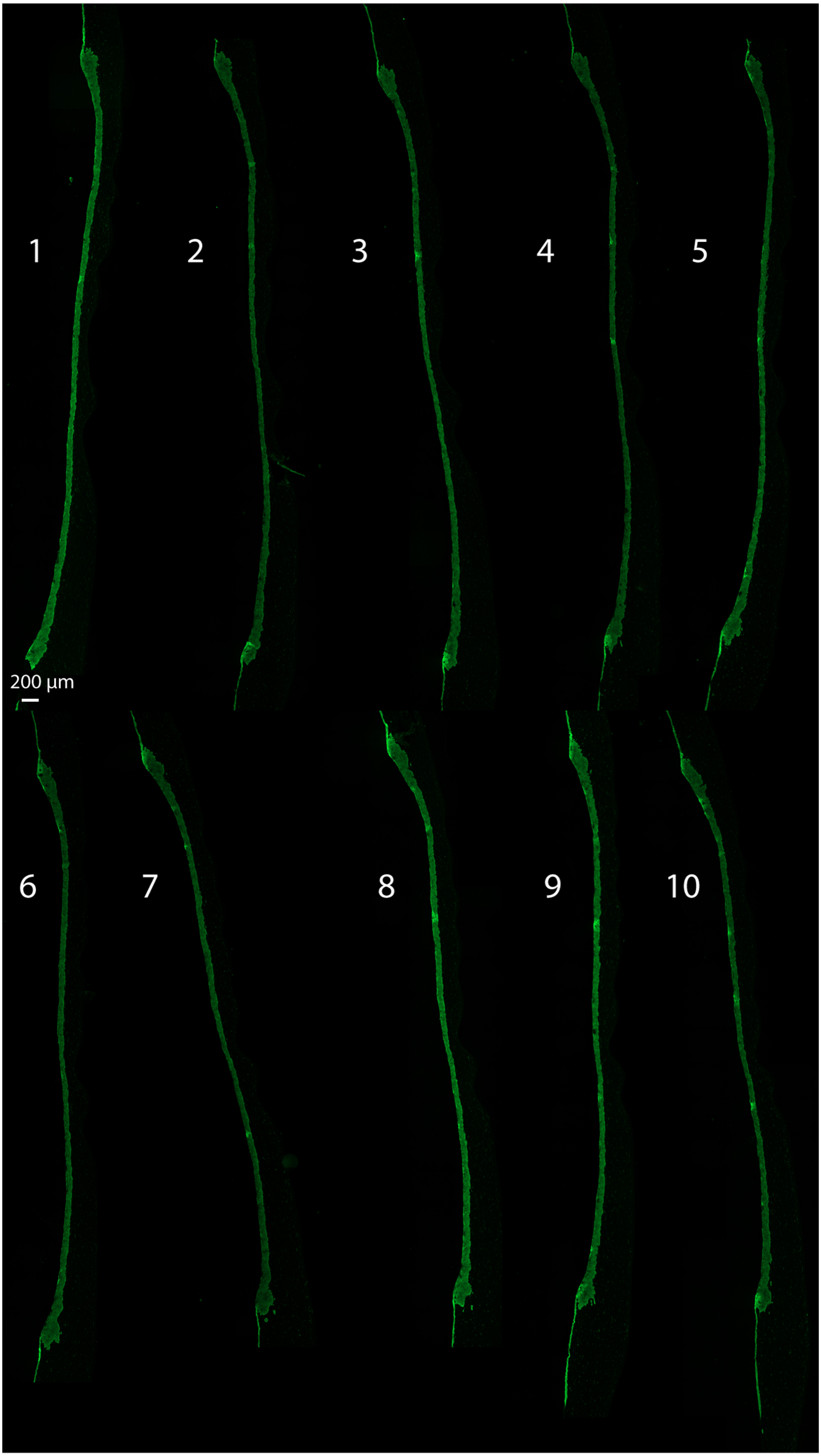

X01105I *C1GALT1* KO 2E5 + HSV-1 36 h section series 2

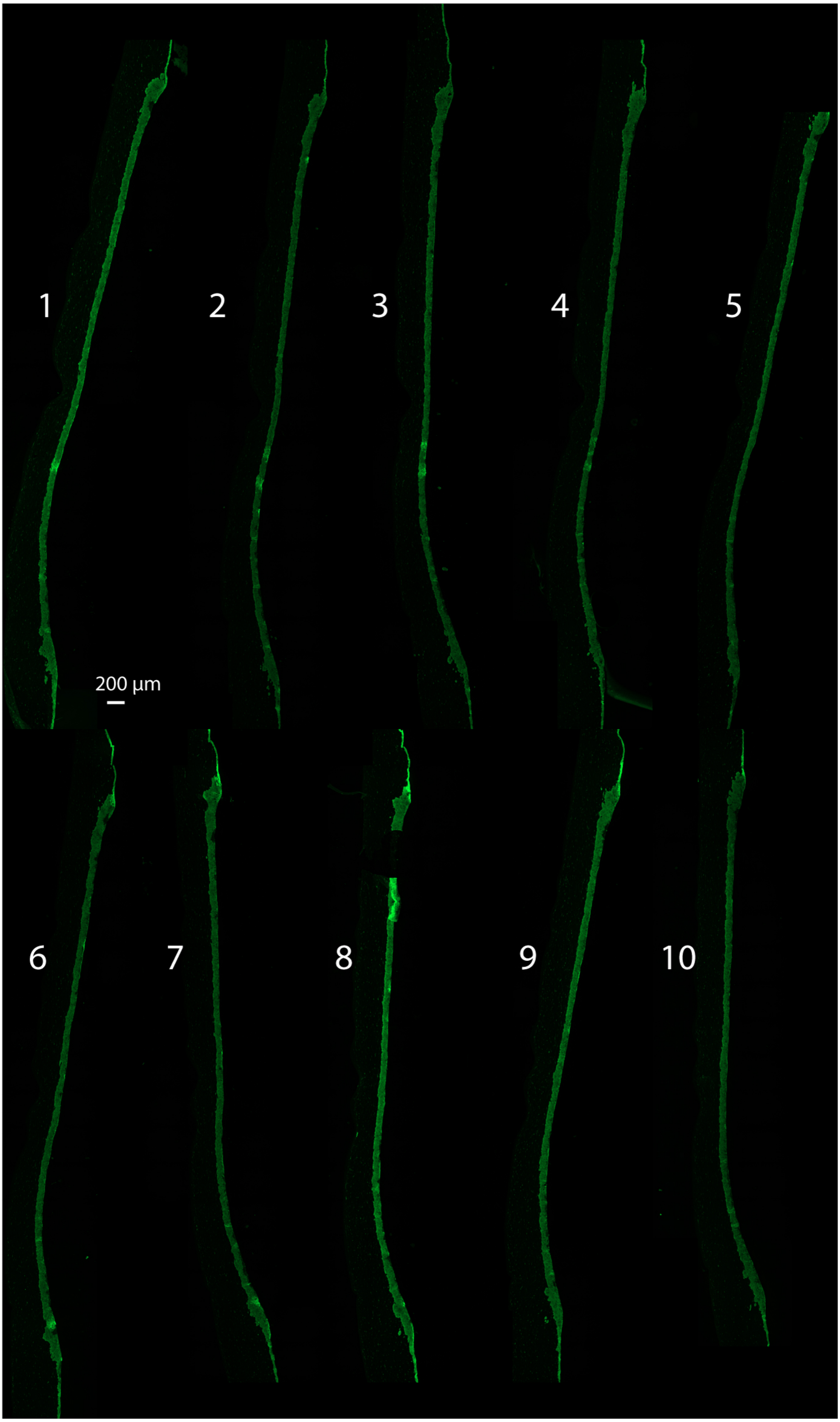

X01105Q *GCNT1* KO 2E7 + HSV-1 36 h section series 1

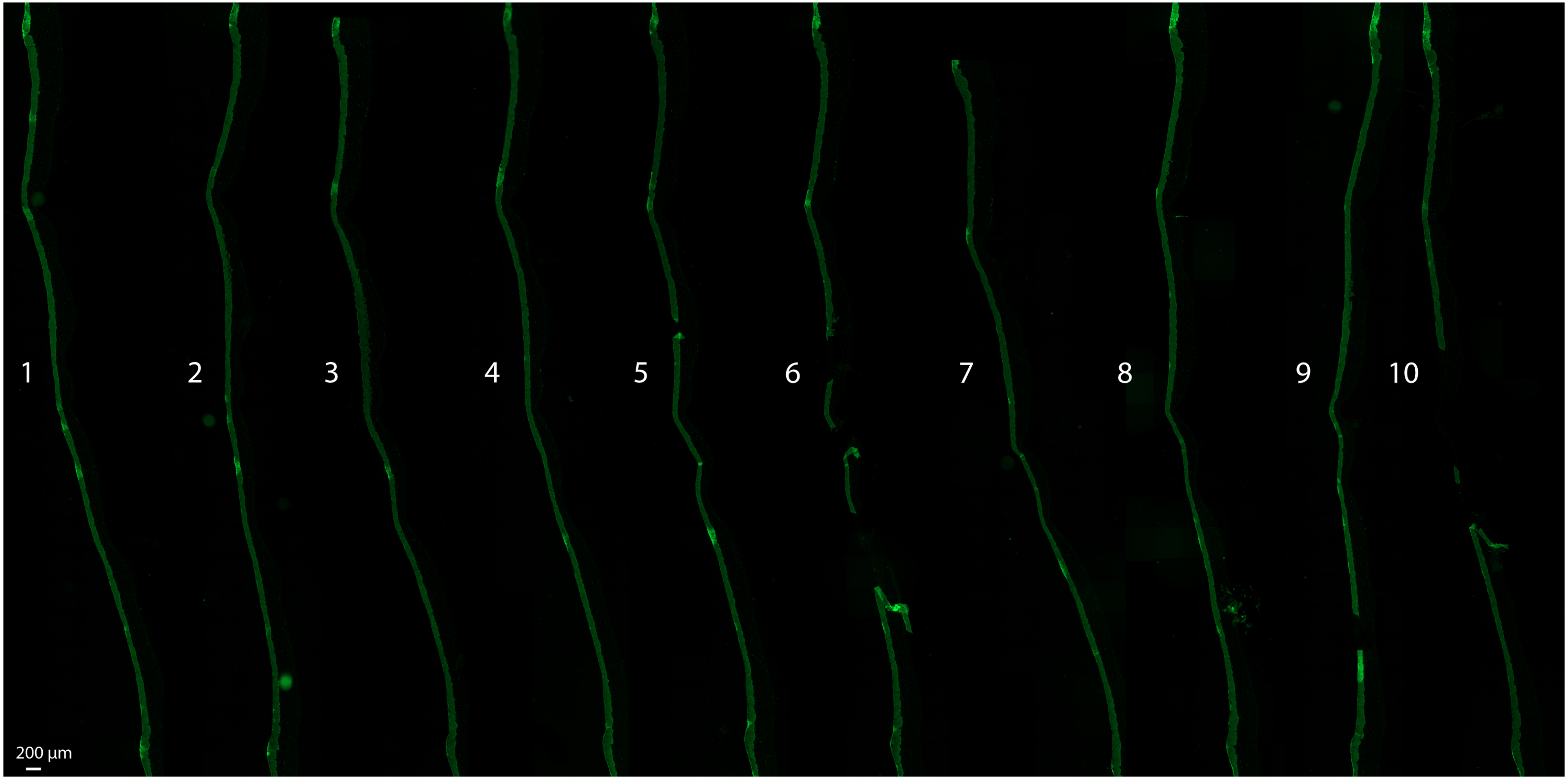

X01105Q *GCNT1* KO 2E7 + HSV-1 36 h section series 2

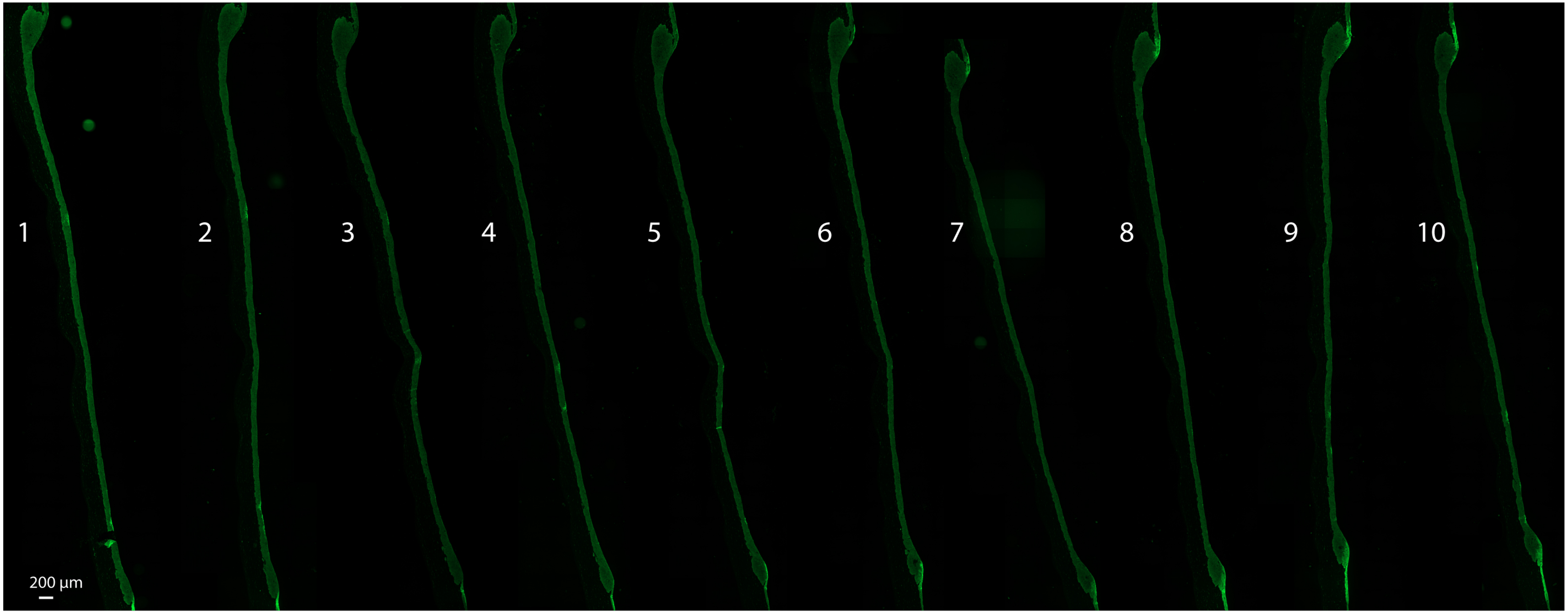

X01105K *ST3GAL1* KO 2B2 + HSV-1 36 h section series 1

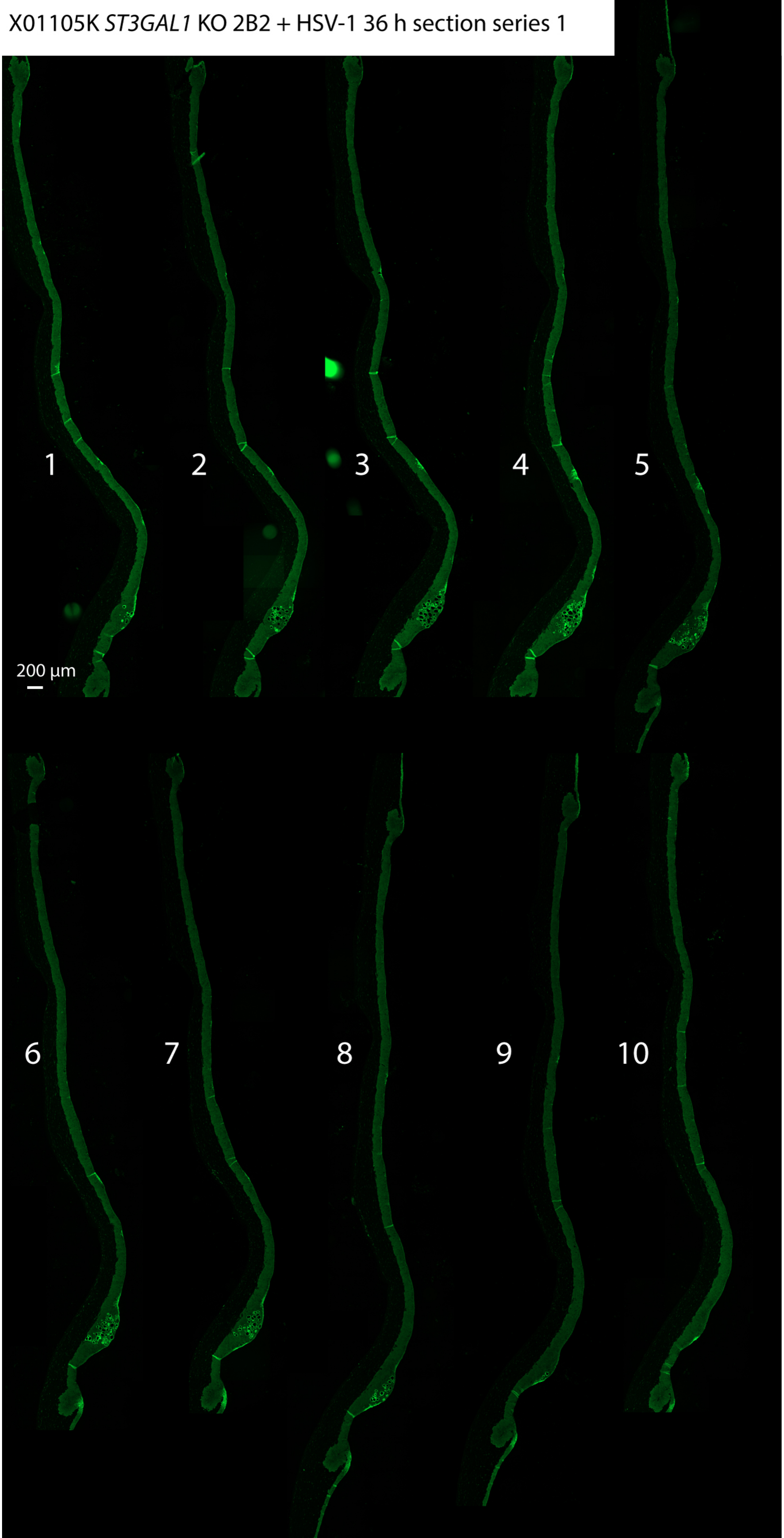

X01105K *ST3GAL1* KO 2B2 + HSV-1 36 h section series 2

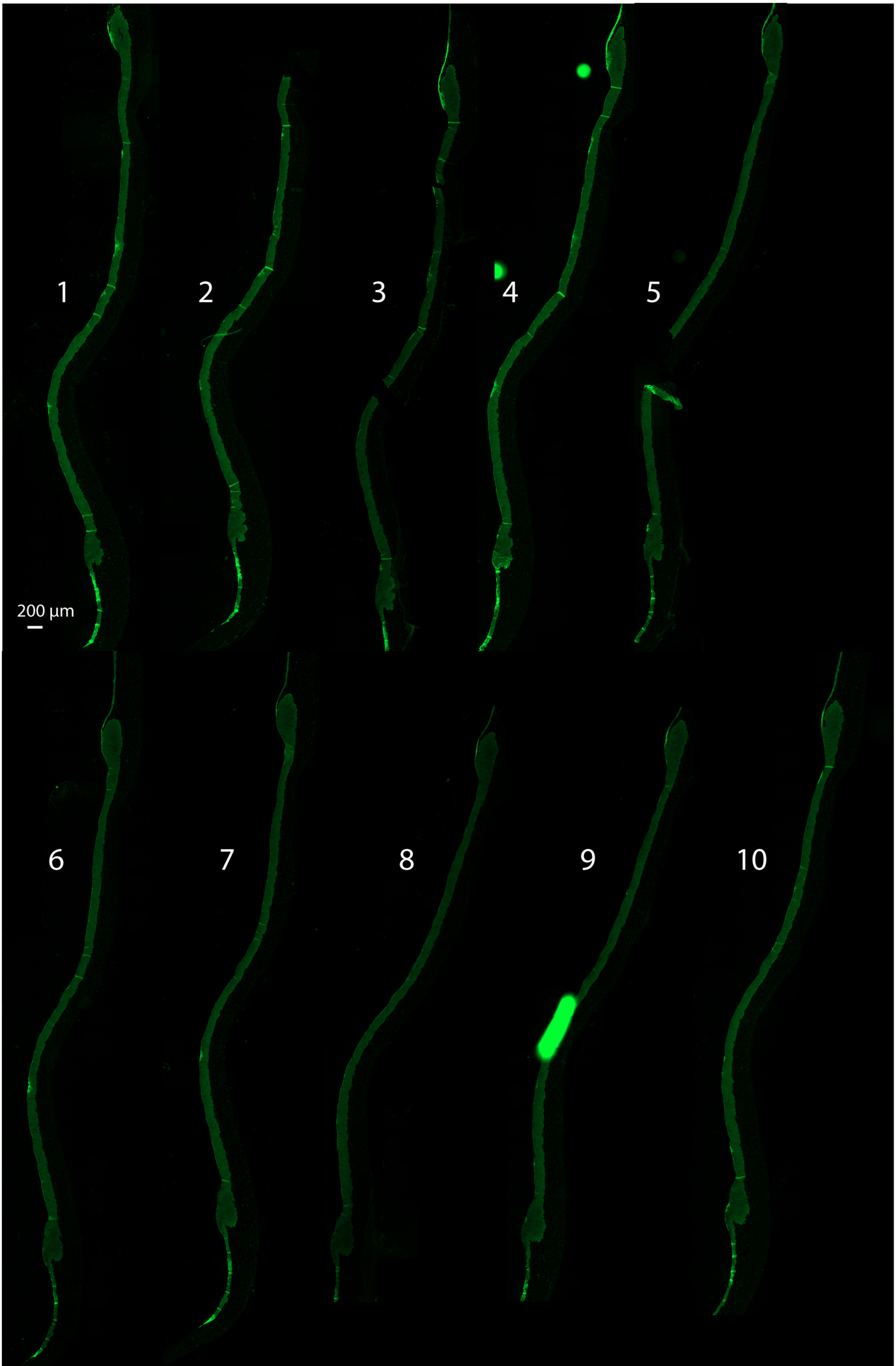

X01105H *B4GALT7* KO 3D9 + HSV-1 36 h section series 1

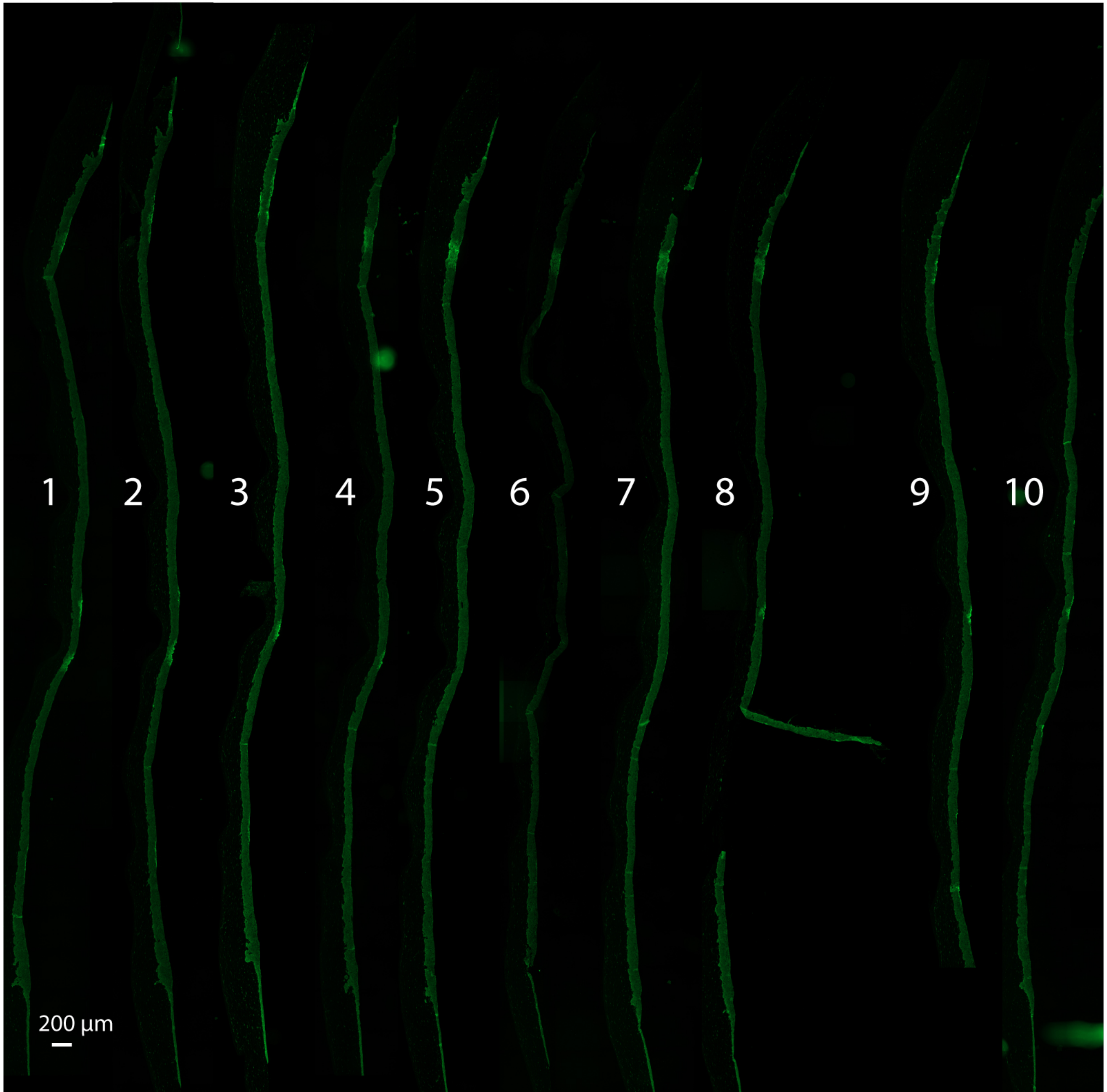

X01105H *B4GALT7* KO 3D9 + HSV-1 36 h section series 2

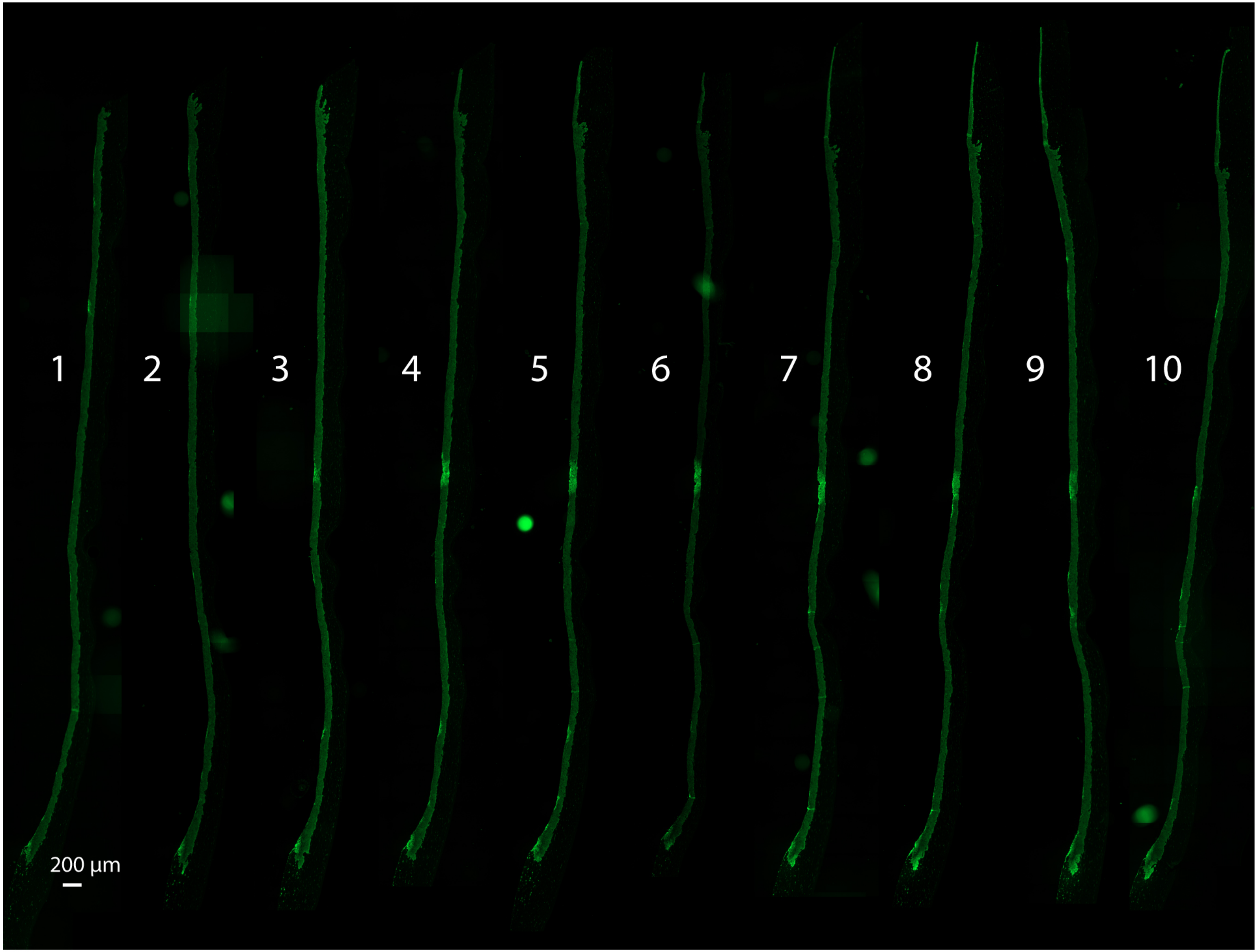

X01105J *B4GALT5* KO D11 + HSV-1 36 h section series 1

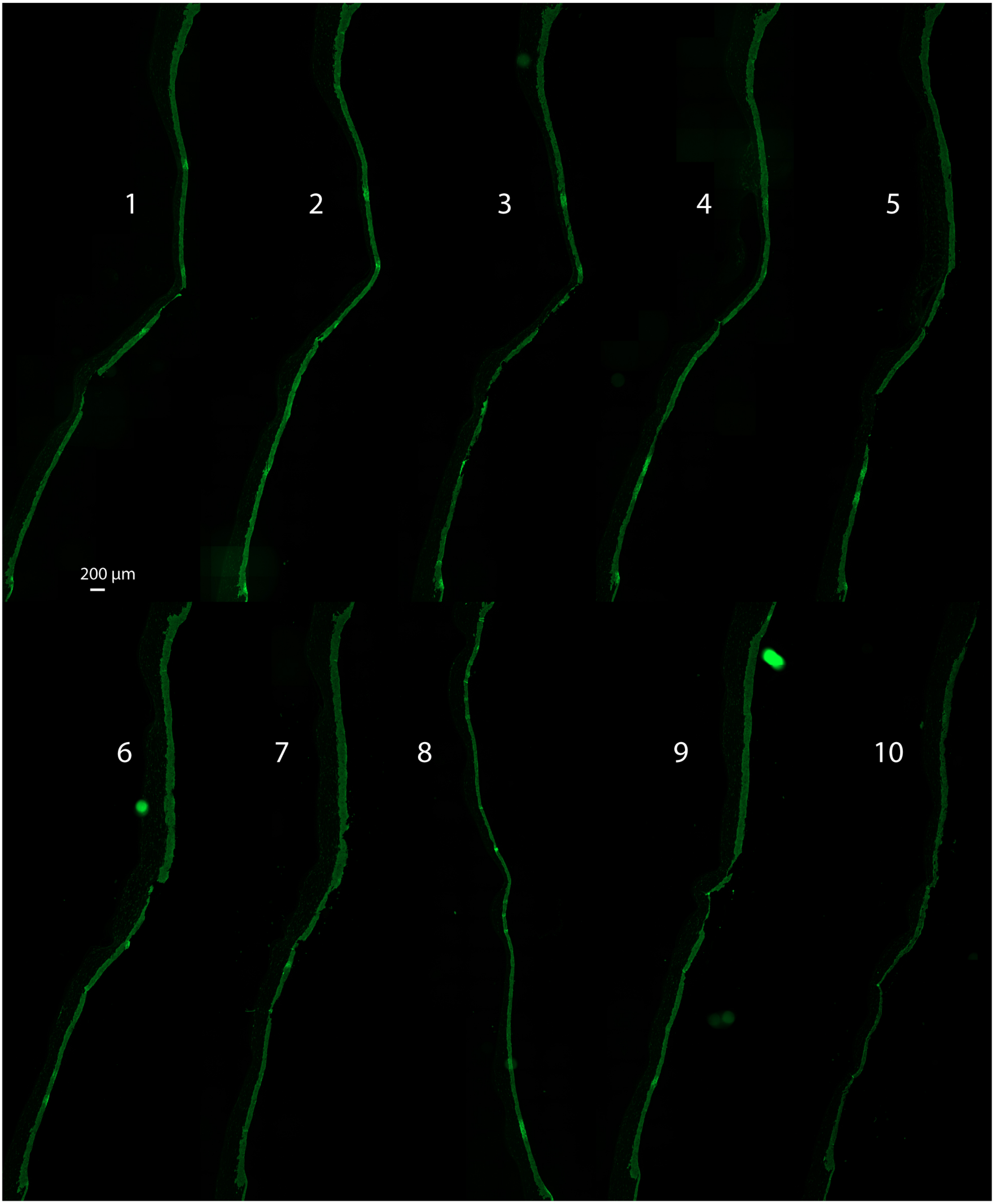

X01105J *B4GALT5* KO D11 + HSV-1 36 h section series 2

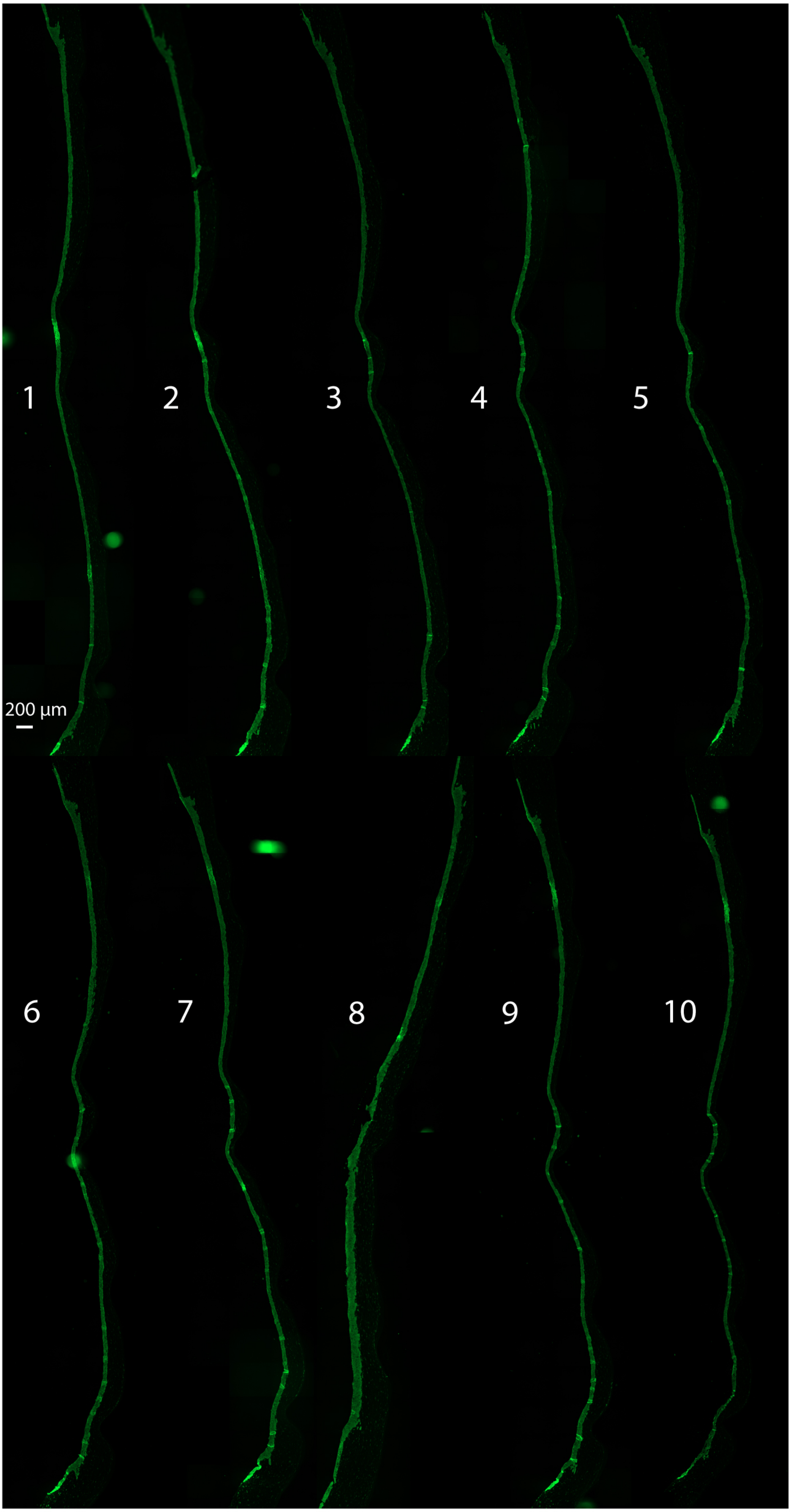

X01105P *ST3GAL5* KO 1C5 + HSV-1 36 h section series 1

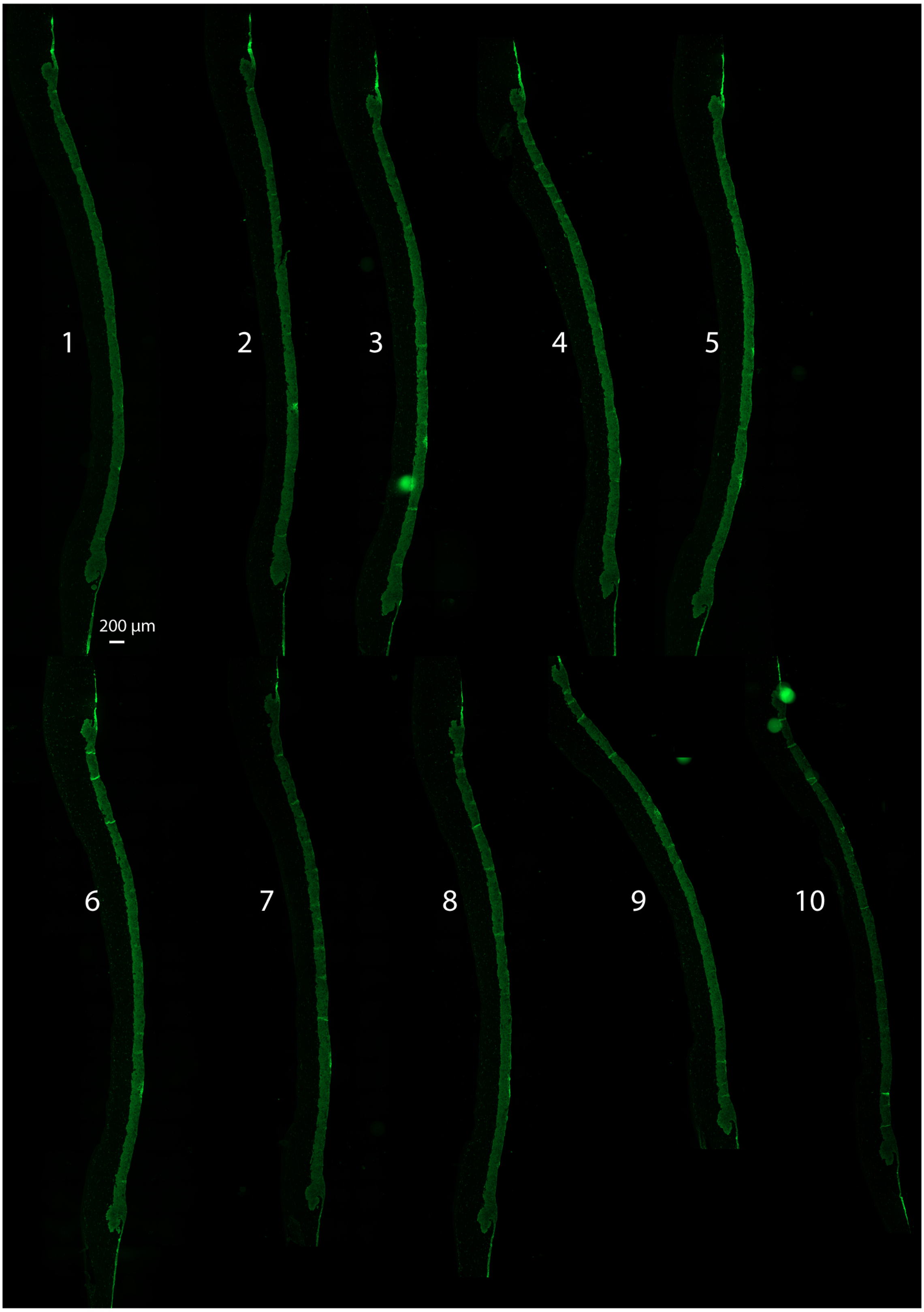

X01105P *ST3GAL5* KO 1C5 + HSV-1 36 h section series 2

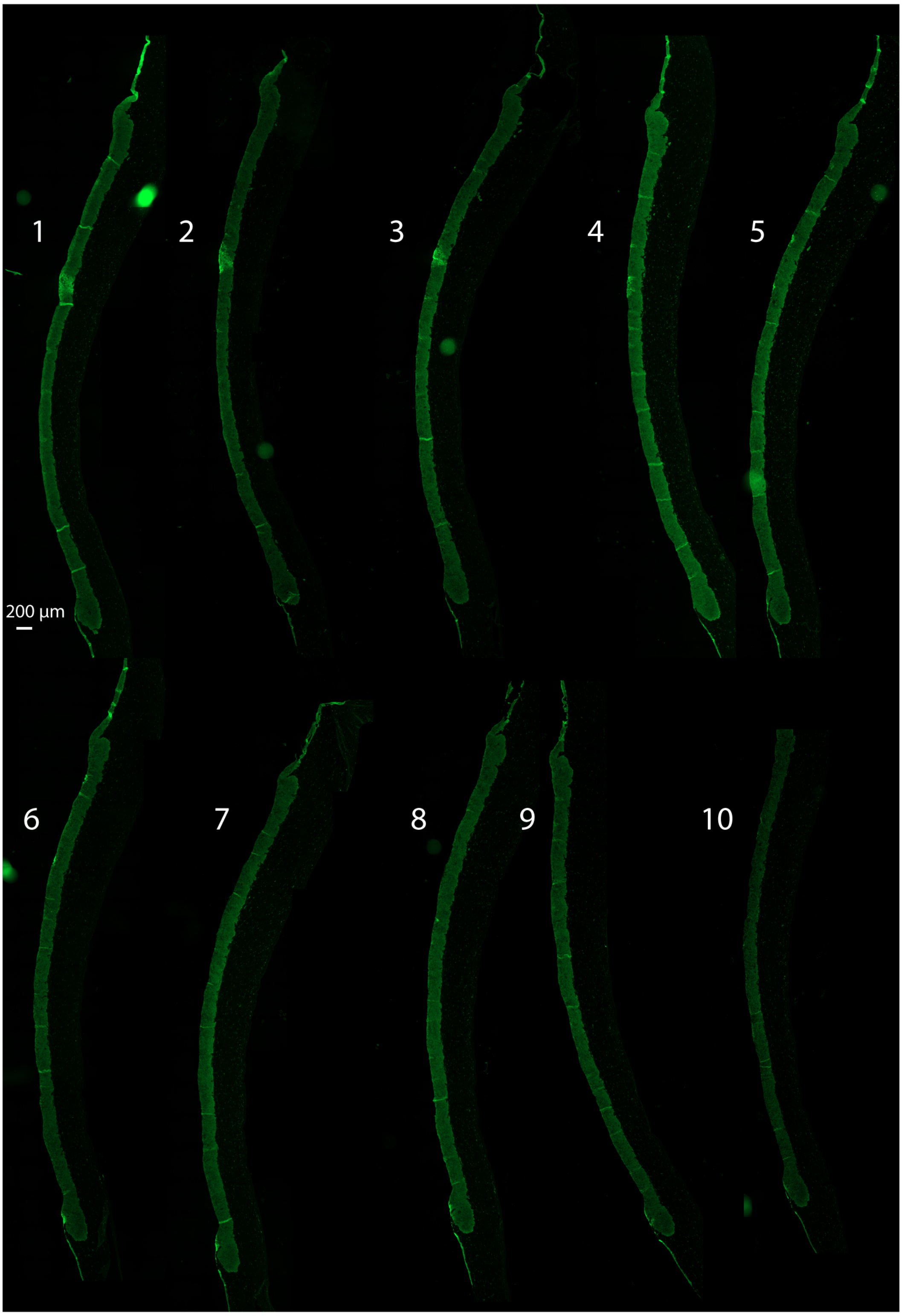
